# Ear pinnae in a neotropical katydid (Orthoptera: Tettigoniidae) function as ultrasound guides for bat detection

**DOI:** 10.1101/2021.09.01.458595

**Authors:** Christian Pulver, Emine Celiker, Charlie Woodrow, Inga Geipel, Carl Soulsbury, Darron A. Cullen, Stephen M Rogers, Daniel Veitch, Fernando Montealegre-Z

**Affiliations:** University of Lincoln, School of Life Sciences, Joseph Banks Laboratories, Green Lane, Lincoln, UK, LN6 7DLf; Smithsonian Tropical Research Institute, Apartado 0843-03092, Balboa, Panama; CoSys Lab, Faculty of Applied Engineering, University of Antwerp, 2020 Antwerpen, Belgium; Flanders Make Strategic Research Centre, 3920 Lommel, Belgium

**Keywords:** bushcricket, bioacoustics, ultrasound hearing, 3D printing, bat predation

## Abstract

Early predator detection is a key component of the predator-prey arms race, and has driven the evolution of multiple animal hearing systems. Katydids (Insecta) have sophisticated ears, each consisting of paired tympana on each foreleg that receive sound directly externally, and internally via a narrowing ear canal through the acoustic spiracle. These ears are pressure-time difference receivers capable of sensitive and accurate directional hearing across a wide frequency range, despite the small size of katydids. Many katydid species have cuticular pinnae which form cavities around the outer tympanal surfaces, but their function is unknown. We investigated pinnal function in the katydid *Copiphora gorgonensis* by combining experimental biophysics and numerical modelling using 3D ear geometries. Results show that the pinnae in *C. gorgonensis* do not assist in directional hearing for specific call frequencies, but instead pinnae act as ultrasound detector devices. Pinnae induced large sound pressure gains that enhanced sound detection at high ultrasonic frequencies (> 60 kHz), matching the echolocation range of co-occurring insectivorous bats. Comparing pinnal mechanics of sympatric katydid species supports these findings, and suggests that pinnae evolved in katydids primarily for enhanced predator detection. Audiograms (both behavioural and neural) and tympanal cavity resonances obtained from living specimens corroborate our findings.

## Introduction

Throughout the animal kingdom, the need to localize sound signals, both to detect conspecifics and to avoid predation, is a major selection pressure. As a result, vastly different species have convergently evolved mechanisms of hearing to fulfil similar functions (Göpfert and Hennig, 2016; Robert, 2005; Warren and Nowotny, 2021), and hearing organs have evolved in closely related taxonomic groups many times independently (Göpfert and Hennig, 2016; Yack, 2004). To determine the location of a sound source, an animal with two ears will utilize interaural time and amplitude differences. Such binaural auditory systems must satisfy three requirements to function: 1) the distance between each ear must be sufficient to produce recognisable differences in sound arrival time, 2) the ears must be separated by a body which is large enough to attenuate sound between them, 3) the ears must be neurologically coupled in order to calculate time and amplitude differences (Lakes-Harlan and Scherberich, 2015; Suga, 1989). However, small animals, such as insects, are too small to exploit diffractive effects of sound on their bodies to perceive minute differences in sound delays and intensities (Michelsen and Larsen, 2008). Katydids (Orthoptera: Tettigoniidae), a family with ∼8,000 species (Cigliano et al., 2021), have overcome this problem by evolving independently functioning ears in their forelegs (Bailey, 1990), thereby increasing the interaural distance. Many species have also evolved the ability to produce and detect ultrasonic frequencies (Strauß et al., 2014), meaning that the resulting distance between the ears provides sufficient spatial separation to exceed the shorter wavelengths of incoming conspecific sounds (Robert, 2005). Each ear receives sound directly at the external tympanal surface, but also internally in a process similar to that of the mammalian ear. In this internal process, sound enters an air-filled ear canal (EC, also called the acoustic trachea) through a specialised opening in the prothorax known as the acoustic spiracle (Kalmring et al., 2003). The EC’s narrowing, exponential horn shape amplifies the sound signals (Celiker et al., 2020a; Michelsen et al., 1994; Veitch et al., 2021) and reduces propagation velocity (Jonsson et al., 2016; Michelsen et al., 1994; Veitch et al., 2021), and leads this decelerated sound signal through the thorax and foreleg to the internal tympanal surface. The combined internal and external inputs means that the tettigoniid ear functions as a pressure – time difference receiver (Michelsen and Larsen, 2008; Robert, 2005; Veitch et al., 2021), unlike the mammalian ear which functions as a single input pressure receiver via the EC.

At the external auditory input, some tettigoniids possess cuticular pinnae (also referred to as folds, flaps or tympanal covers) partially enclosing their tympana (Fig. 1*A-C*). The role(s) of these pinnae are unclear. Early observations by Autrum suggested that pinnae aid the insect to determine the direction of sound, effectively acting as a sound guide (Autrum, 1963, 1940). The prominence of cuticular pinnae present in a variety of Pseudophyllinae and Conocephalinae species (see examples in fig. S1*A, C-K*) generated more interest in these observations.

**Figure 1.**
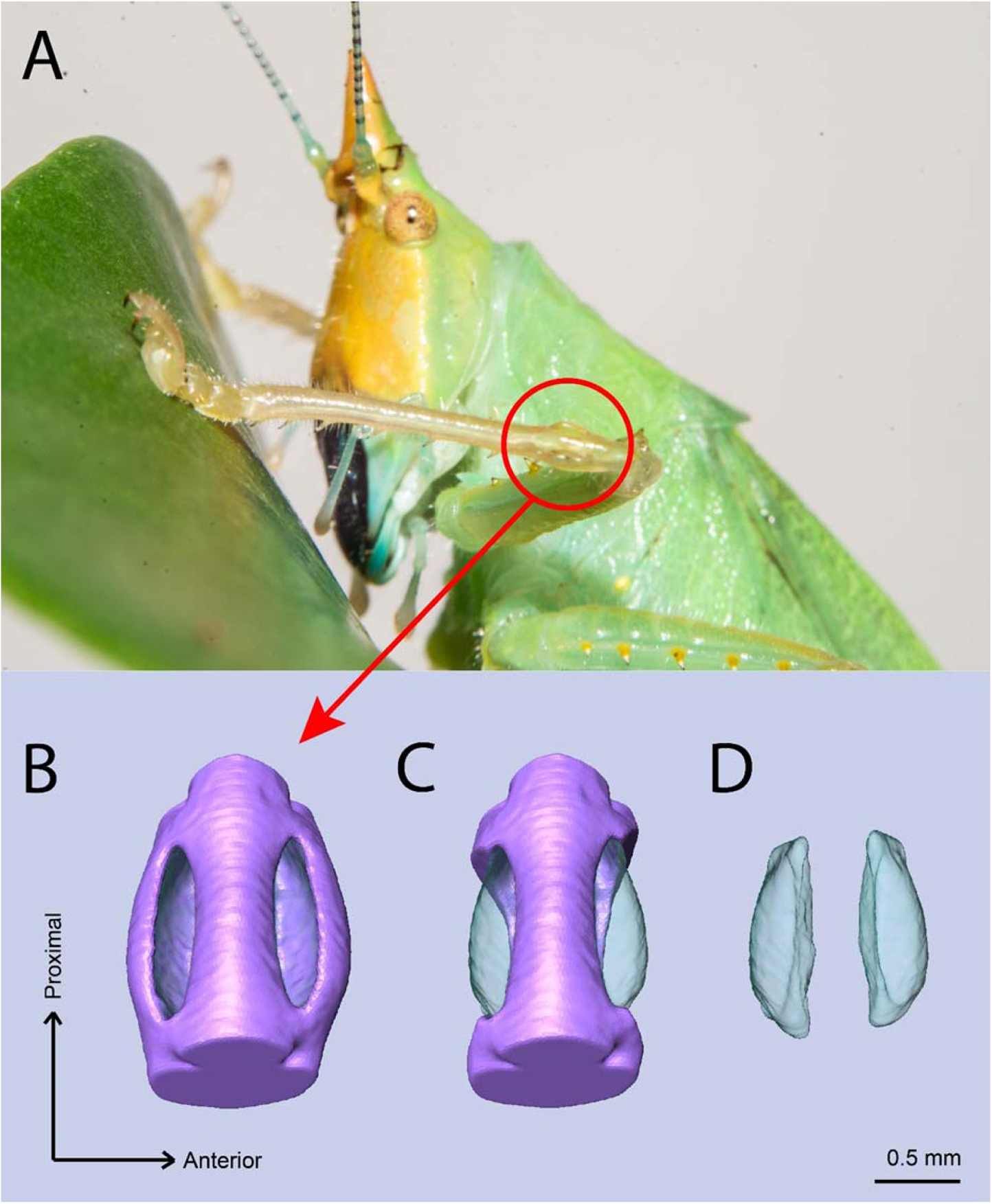
The ear of *Copiphora gorgonensis.* (**A**) Location of the ear in the foreleg. (**B**) 3D anatomy of the ear, with pinnae present; (**C**) 3D anatomy of the following pinnae ablation, with the volume of the subslit cavities exhibited (light blue); (**D**) 3D model of only the subslit cavities.

Morphologies of cuticular pinnae vary greatly between species, and were originally categorized in a phylogenetic context (Bailey, 1990). A dual functioning auditory system in the tettigoniid *Mygalopsis marki* was proposed to explain differences in the auditory morphologies in spiracle size and tympanal pinnae in which the external tympanal ports appear to function as omnidirectional receivers and the EC combined with the spiracle operate as a highly sensitive non-directional receiver (Bailey and Stephen, 1978). These findings were corroborated in *Hemisaga* sp. where an increased acoustic sensitivity at the external tympanal port (through blocking the entrances or slits) was demonstrated (Stephen and Bailey, 1982). A dual channel system consisting of the EC and spiracle serve for the detection of predators, whilst the external tympanal ports are used for detecting conspecific communication signals (Stephen and Bailey, 1982). Studies of ultrasonic rainforest Pseudophyllinae provided more evidence of principal sound reception for conspecific communication using the external tympanal port as a consequence of exceptionally small spiracle sizes (Mason et al., 1991). It was reported that diffraction of very short wavelengths along the tympanal slits contributed to directional orientation in rainforest katydids (Mason et al., 1991).

Before experimental evidence of the dual port system in katydids was published (Jonsson et al., 2016; Michelsen et al., 1994), attention was given to the EC as the main port for sound capture (Eisner and Popov, 1978; Hill and Oldfield, 1981; Hoffmann and Jatho, 1995; Michelsen and Nocke, 1974; Shen, 1993). Moreover, it has been argued that the size of the ear is considerably smaller than the wavelengths of most carrier frequencies of described insects known at the time, and therefore the sound pressure field around the ear would be constant and yield no directional cues (Michelsen and Nocke, 1974). In other words, sound accessing the external tympanal port is not related to the direction of incidence, and tympanal pinnae are merely protective features sheltering the fragile tympanum. However, recent research showed that quieter, low amplitude sound waves acting on the external tympanal membrane (without gain from the EC) of the katydid *Copiphora gorgonensis* (Tettigoniidae: Copiphorini) do cause vibrations of significant amplitude in the inner ear (Montealegre-Z and Robert, 2015). Therefore, even these very weak vibrations are mechanically transduced. The external sound arrives 60-80 µs before its amplified form of self on the internal tympanal surface via the EC, and this significant phase delay forms the basis of the pressure – time difference receiver definition (Jonsson et al., 2016; Veitch et al., 2021). In katydids with cuticular pinnae surrounding the tympana, evidence suggests that the insect can use both ports, but how the external port contributes to directionality remains unknown (Römer, 2020).

Here, we investigate the role of cuticular pinnae using the neotropical katydid *Copiphora gorgonensis*, a species endemic to Gorgona, an island in the Pacific Ocean off the western coast of Colombia (Montealegre-Z and Postles, 2010). Males of this species produce a pure-tone song a 23 kHz to attract females. *Copiphora gorgonensis* has become a model species for hearing studies due to the transparency of the cuticle which facilitates non-invasive, real-time measurements of the inner ear (Montealegre-Z et al., 2012; Montealegre-Z and Robert, 2015).

We integrated experimental biophysical measurements based on micro–scanning laser Doppler vibrometry (LDV) and micro-computed tomography (µ-CT) to simulate the function of the cuticular pinnae and how they contribute to auditory orientation in this katydid. The pairing of these approaches were applied to 3-dimensional (3D) print models of the ear, and scaled experiments were performed to validate the simulations. We investigated if: 1) the direction of incidence of the sound stimulus, presented by a loudspeaker, is a function of the sound wave directly accessing the tympana through the slits; 2) the tympanal cavities produce sound pressure gains that act externally on the tympana; 3) tuning properties of the pinnal cavities are a result of pinnal geometry and can be predicted by the volume and/or entrance size of the cavity.

Based on Autrum’s original observations, we hypothesized that tympanal pinnae function as ultrasonic guides by pinnae forming exceptionally small resonant cavities. Further, we hypothesized that these cavities act as Helmholtz-like resonators able to capture and amplify diminishing ultra-high frequency sound waves.

## Results

We investigated the role of tympanal pinnae in sound capture by testing how the direction of incidence of the sound stimulus presented by the loudspeaker induced tympanal displacements at three frequencies (23, 40 and 60 kHz) with the cuticular pinnae intact and later ablated. Frequencies above 60 kHz were not tested provided the acoustic limitations of the experimental platform and the poor performance of the probe speaker at high ultrasounds (see *Materials and Methods*). A total of 2,736 measurements were performed on 13 ears (1,512 measurements for four male specimens; 1,224 for three female specimens). For time of sound arrival, we found a significant interaction between the presence of cuticular pinnae with angle of incidence (21° azimuth frontal to ear; Fig. 2*A*) and with frequency (table S1). Post hoc analysis showed that the presence of pinnae significantly delayed the time of arrival to ipsilateral tympana at 23 kHz (t-ratio = −11.15, *P* < 0.001) and 40 kHz (t-ratio = −7.43, *P* < 0.001), but not at 60 kHz (t-ratio = −1.86, *P* = 0.063). The effect of tympanum (anterior or posterior) on time of arrival and displacement amplitudes was found to be significant with the PTM exhibiting slightly longer arrival times (∼ 2 µs) but a 21.5% lower mean displacement amplitude (Fig. 2*B*; table S1).

**Figure 2.**
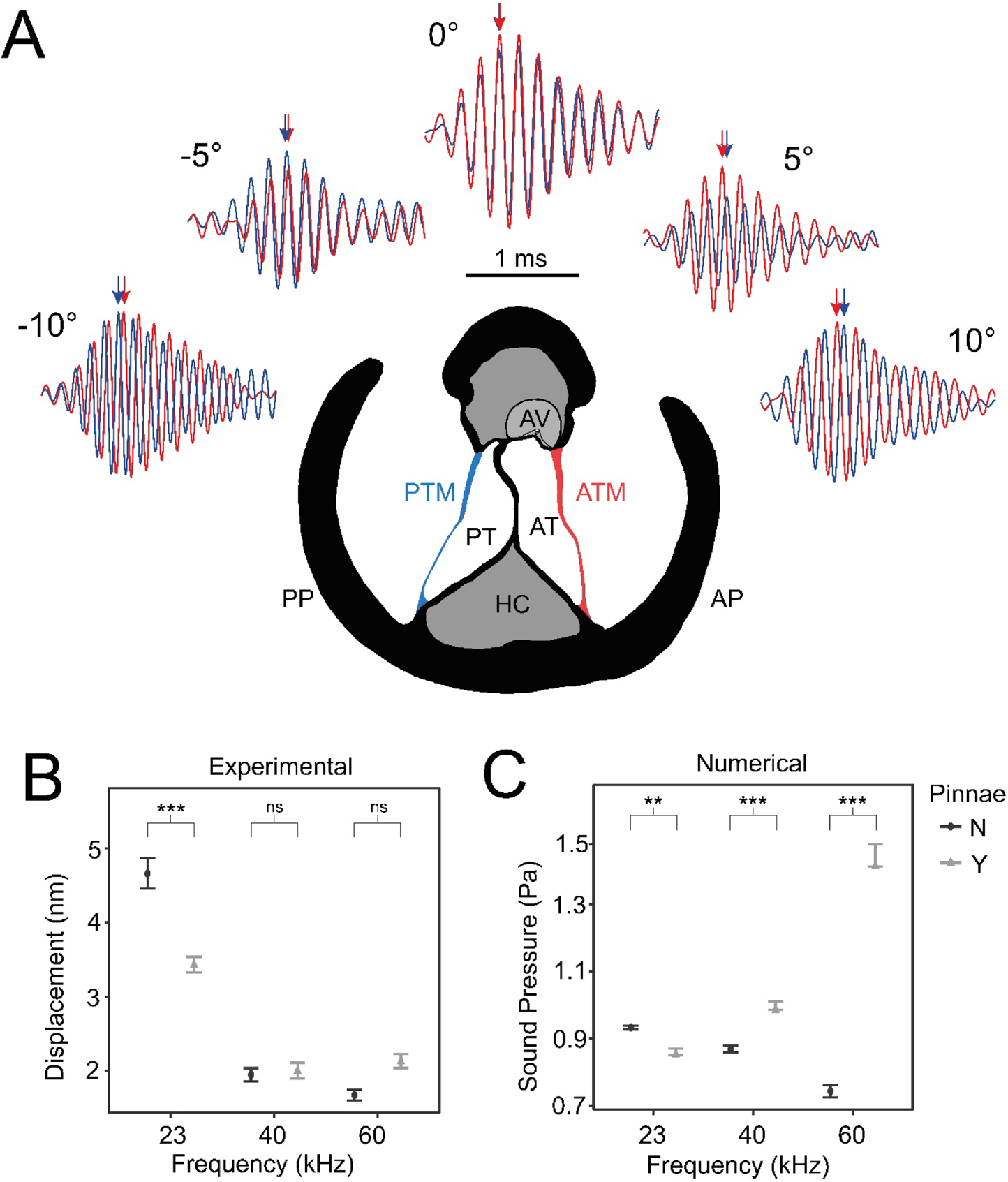
The effect of pinnae in the time domain and numerical simulations. (**A**) Time plots from five incidence angles for the 60 kHz test sound illustrating changes in oscillation phase between the anterior (ATM, in red) and posterior (PTM, in blue) tympana of the same ear. Notice the phase difference of 0.25 cycles is 90° at −10° and 10°. An anatomical cross section of the ear is shown with each tympanum (ATM and PTM), auditory vesicle (AV), posterior and anterior bifurcated tracheal branches (PT and AT), haemolymph channel (HC) and posterior and anterior pinnal structures (PP and AP). (**B**) Mean displacement amplitudes (nm) of the tympanic membranes for each tested frequency (23, 40 and 60 kHz) with and without the presence of cuticular pinnae (*n* = 9 ears). (**C)** Cavity induced pressure gains with pinnae compared to sound pressure (Pa) predictions with the pinnae ablated from numerical models using Comsol Multiphysics (17 ears; 10 females, 7 males). For means comparison plots (**B** & **C**), significance symbols from post hoc analyses: ‘***’ 0.001, ‘**’ 0.01, ‘*’ 0.05, ‘ns’ 0.1, and ‘ ’ 1. grey bars with cuticular pinnae and black bars without cuticular pinnae showing standard error.

For displacement amplitude, there was a significant interaction between the presence of pinnae and frequency (table S1). Post hoc analysis showed greatest displacement amplitudes at 23 kHz with the pinnae ablated (t-ratio = 3.20, *P* < 0.001; Fig. 2*B*). This demonstrates that pinnae do not enhance auditory perception of the carrier frequency in *C. gorgonensis*, and that the tympanum achieves this displacement by resonance (see (Jonsson et al., 2016; Montealegre-Z et al., 2012)). There were no differences in displacement at either 40 kHz (t-ratio = 0.84, *P* = 0.399; Fig. 2*B*) and 60 kHz (t-ratio = −0.61, *P*=0.540; Fig. 2*B*) regardless of pinnal condition. We also found a significant interaction between the presence of cuticular pinnae with angle of incidence, showing the impact of pinnae in increasing arrival time as the sound source rotates opposite the ear (table S1). Responses were strongest for sound presented perpendicular to each respective slit cavity (average 3.08 ± 2.91 nm at 10°, average 2.90 ± 2.98 nm at −9°) with the lowest displacement amplitudes occurring when sound was directed at the region of the dorsal cuticle bifurcating the cavities, also referred as “point zero”, showing (average 1.99 ± 1.90 nm at −1°) with the pinnae intact due to cuticle obstructing the response of the tympanal membrane. In contrast, point zero and adjacent angles showed the greatest displacement amplitude with the pinnae ablated (average 3.04 ± 3.42 nm at −1°) with incident angles on either side of point zero showing a gradual subdued response to the stimulus (average 2.73 ± 3.27 nm at 10°, average 2.54 ± 2.57 nm at −10°).

Phase angle (φ*°*) was calculated from the absolute value of the difference between the vibrations of the ATM and PTM per recording (1,532 in total). Pinnae maintained mean Δφ*°* at 80.9° for 23 kHz, 88.8° for 40 kHz, and 84.1° for 60 kHz, but with the pinnae ablated, phase differences were smaller particularly at 60 kHz (Δφ*°* at 62.7° for 23 kHz, 78.7° for 40 kHz, and 49° for 60 kHz).

### Anatomical measurements of the external tympanal port

The anatomical features of the ear were measured to predict resonance and compare intraspecific variation in pinnal size (table S2). 2D measurements of the area of the pinnal opening (slit), distance between the centre of the ear (septum) and edge of the pinna (pinnal protrusion), and distance between slits (septum width) were studied using an Alicona Infinite Focus microscope. 3D measurements of the cavities and cross section of the ears were performed with the µ-CT scanner using the software Amira-Aviso 6.7 (*n* = 8 ears; 3 females and 2 males). We found that the average size of the slits (0.16 ± 0.01 mm^2^) and cavities (0.14 ± 0.01 mm^3^) were nearly identical between the ATM and PTM. The pinnal protrusion length showed that the PTM pinnae (0.44 ± 0.03 mm) were wider than the ATM (0.39 ± 0.02 mm). The mean cross-sectional width of the ear was 1.14 ± 0.35 mm.

### Tympanal cavity resonance calculations

We used the 2D (slit area) and 3D measurements (cavity volume) to estimate resonance of the tympanal cavities (table S2). This was calculated with the assumption that the slit openings are a perfect circle (to determine radius) and the cavity acts as a cylindrical tube using a neckless Helmholtz resonance equation. Here, *c* is speed of sound in air (343 m s^−1^), cross-sectional area of the opening with radius *r*, 1.85 is the correction length of the neck and *V* denotes the volume of the resonator/cavity (Rossing and Fletcher, 2004).

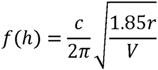

The pinnal cavities (*n* = 8) showed a neckless Helmholtz resonance of 94.28 ± 3.53 kHz for the ATM and 91.69 ± 3.93 kHz for the PTM. These calculations suggest that the pinnae cavity does not resonate at the specific calling song frequency of *C. gorgonensis*.

### 3D printed model time and frequency domain measurements of pinnal cavities

3D printed scaled models of the ear were used to measure sound pressure gains and resonances, to overcome the problem of experimental limitations imposed by the small size of the specimens and ears. 3D ears (*n* = 8; 1 male and 1 female ± pinnae) were printed at a scale of 1:11.43 and the acoustic stimuli was scaled by the same factor for pure tones (2.01 kHz for 23 kHz, 3.50 kHz for 40 kHz, 5.25 kHz for 60 kHz, and 9.63 kHz for 110 kHz) and for broadband (2 to 15 kHz for 11.5 to 170 kHz). The interaction between pinnae and frequency significantly affected sound pressure (dB), whilst tympanum (ATM or PTM) did not (table S1). Across all frequencies, the presence of pinnae increased sound pressure, but differences were greatest at higher frequencies (23 kHz: t-ratio = −2.54, *P* = 0.014; 40 kHz: t-ratio = −8.69, *P* < 0.001; 60 kHz t-ratio = −15.66, *P* < 0.001; 110 kHz t-ratio = 41.70, *P* < 0.001; Fig. 3*A*), the greatest pressure gains were detected at 101.47± 3.43 kHz for both the ATM (26.33 ± 4.06 dB) and PTM (30.04 ± 1.34 dB) with the pinnae intact. With the pinnae ablated, the greatest pressure gain was found to be at 101.41 ± 0.86 kHz for both the ATM (9.69 ± 0.87 dB) and PTM (9.83 ± 0.97 dB).

**Figure 3.**
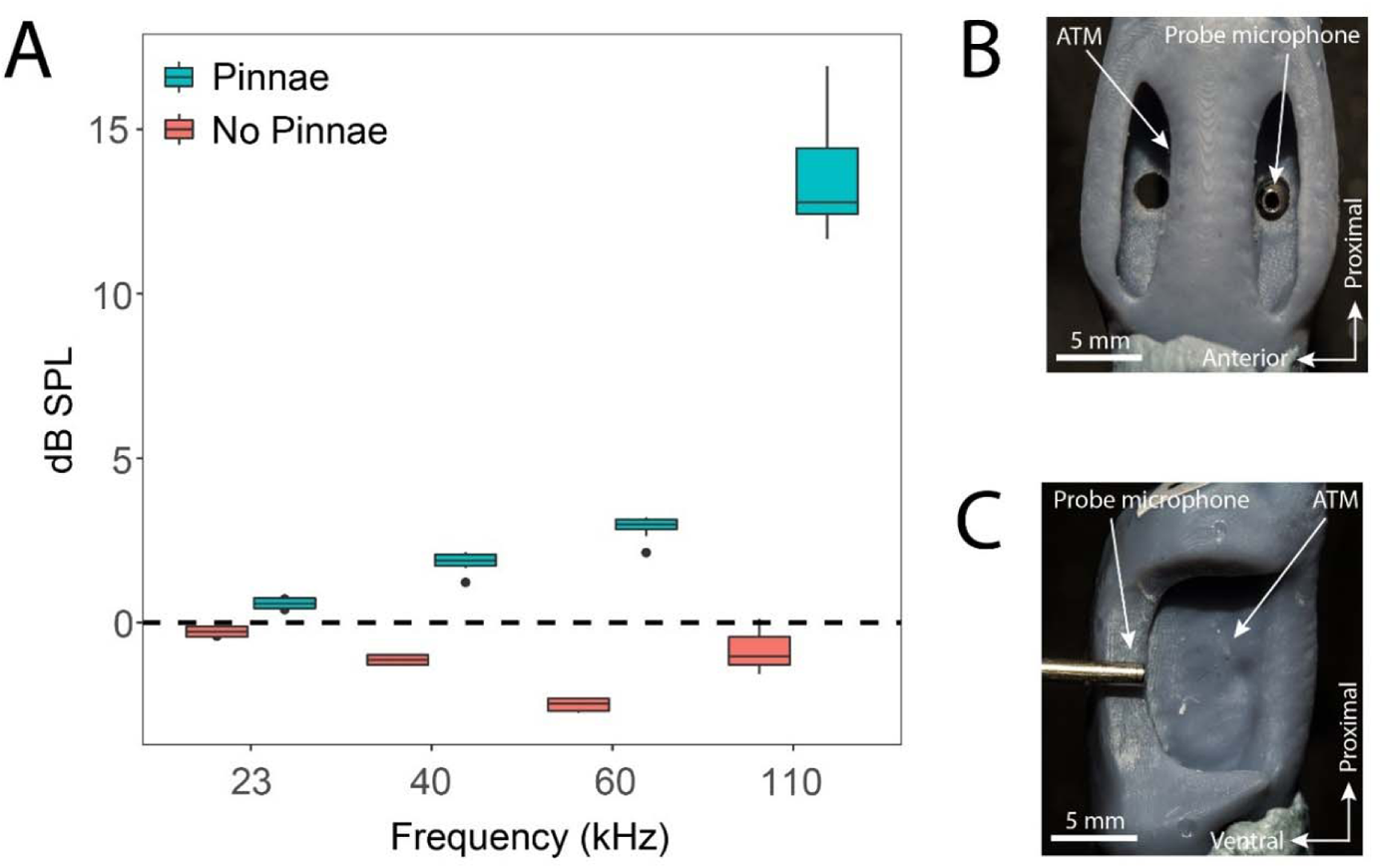
Acoustic experiments with 3D printed scaled ear models. ***A***. Sound pressure gains (dB SPL) of 3D printed ears calculated from scaled time domain recordings for 23, 40, 60 and 110 kHz 4-cycle pure tones. ***B.*** An example of a 3D printed ear model with pinnae present (dorsal view) showing the probe microphone inside the PTM. ***C***. An example of a 3D printed ear model with pinnae ablated (anterior lateral view), showing probe microphone placement.

### Numerical results

Using real 3D geometries of each experimental ear (*n* = 17 ears; 8 with pinnae, 9 with pinnae ablated), we used Finite Element Analysis (FEA) to simulate sound pressure gains and the effect of incident angle at frequencies exceeding the experimental limitations on living specimens (see *Materials and Methods*). For sound pressure measurements there was a significant interaction between the presence of pinnae and frequency (table S1). At 23 kHz ears without pinnae had significantly higher sound pressures (t-ratio = 3.45, *P* < 0.001), but the effect was reversed at 40 kHz (t-ratio = −5.94, *P* < 0.001) and 60 kHz (t-ratio = −28.52, *P* < 0.001), with differences increasing as frequency increased (Fig. 2*C*). There was no effect of angle of sound incidence (−10°, −5°, 0°, 5°, 10°) or tympanum on sound pressures (table S1).

Simulated sound pressure gains (Figs. 4*C* and 4*D*), and their distribution maps (Fig. 4*A* and *B*) showed the greatest sound pressure gain at a mean value of 118 kHz (ATM 121 kHz, PTM 115 kHz), and such gains were reduced or lost entirely when the pinnae were removed (table S1; Fig. 4*D*).

**Figure 4.**
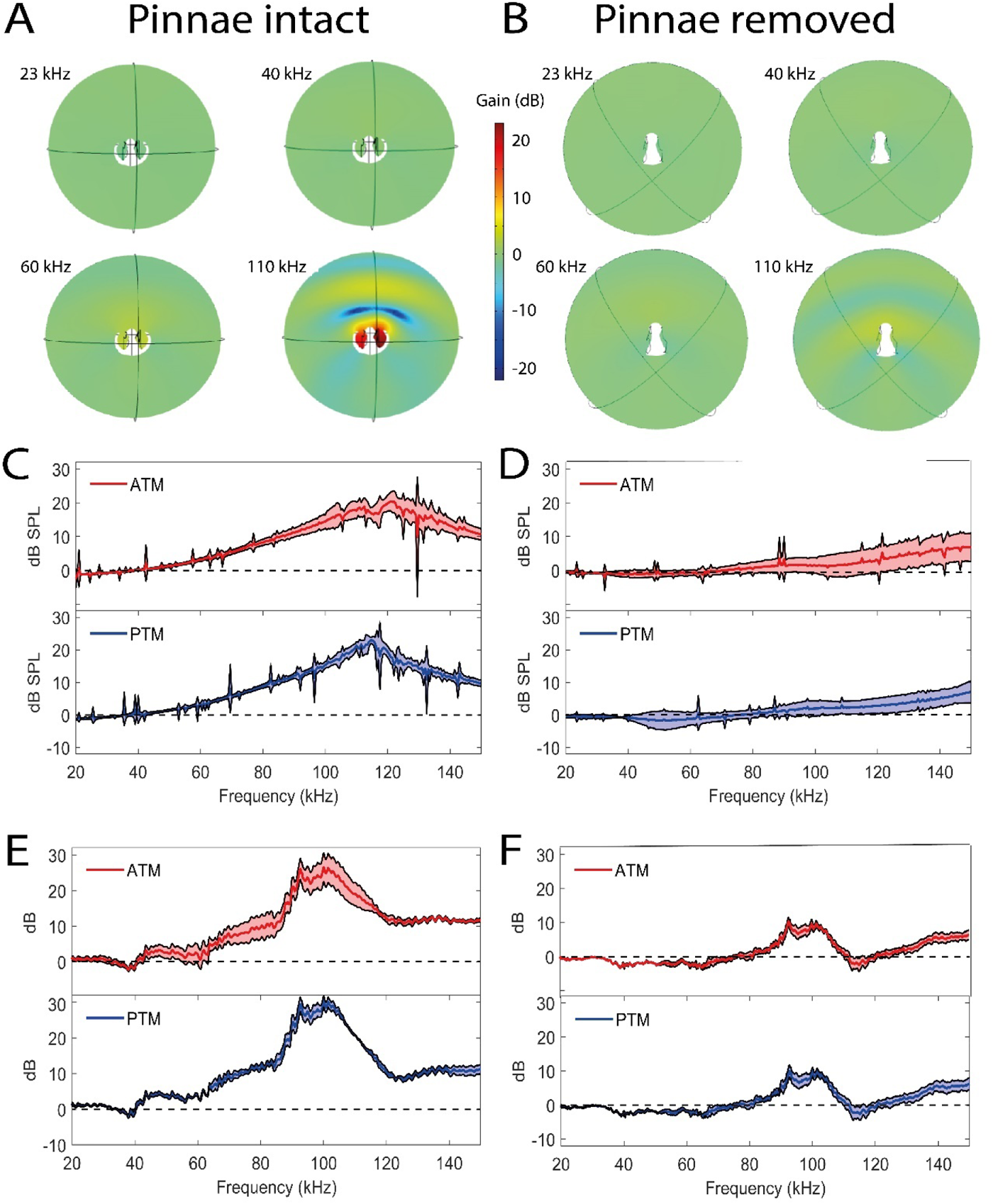
Sound pressure gains measured by numerical simulations of sound capture in the pinnae cavities (using COMSOL Multiphysics) and experimentally using printed 3D-scaled ear geometries. Panels ***A***, ***C***, and ***E*** depict cavity induced sound pressure distribution and gains with pinnae, panels ***B***, ***D***, and ***F*** represent sound pressure gains without the pinnae. ***A*** and ***B***. Cross – section of the ear of *Copiphora gorgonensis* with the pinnal structures intact (***A***) and ablated (***B***). Sound pressure intensities depicted with colours for simulations of 23, 40, 60 and 110 kHz. Low sound pressure dB (blue) to high sound pressure dB (red) distributions inside and outside the cavities. ***C*** and ***D*** are plots of simulated sound pressure gains (dB SPL) in the frequency ranges of 20 to 150 kHz for each tympanum. ***E*** and ***F*** are plots of relative dB gain of the tympanal cavities in the 3D printed ears. ATM in red and PTM in blue.

The effects of angle, pinnae, tympanum, interaction of angle and pinnae, and the interaction of pinnae and frequency were not significant on arrival times. However, frequency was a significant factor on arrival times (23 vs. 40 kHz: t-ratio = 30.739, *P* < 0.001; 23 vs. 60 kHz: t-ratio = 45.857, *P* < 0.001; 40 vs. 60 kHz: t-ratio = 15.117, *P* < 0.001).

### Tympanal and behavioral response to broadband stimulation

For broad tympanal responses, we exposed seven specimens with intact pinnae to broadband periodic chirp stimulation in the range 20 to 120 kHz in a free sound field and recorded the vibration of both ATM and PTM, of the two ears using a micro-scanning laser Doppler vibrometer. There was a relatively stable response (measured as velocity per sound pressure) of the tympanal membranes between 20 to 70 kHz. However, above 80 kHz the tympana response increased dramatically with resonant peaks at 107.84 ± 3.74 kHz for the PTM and 111.13 ± 4.24 kHz for the ATM (Fig. 5*A*).

**Figure 5.**
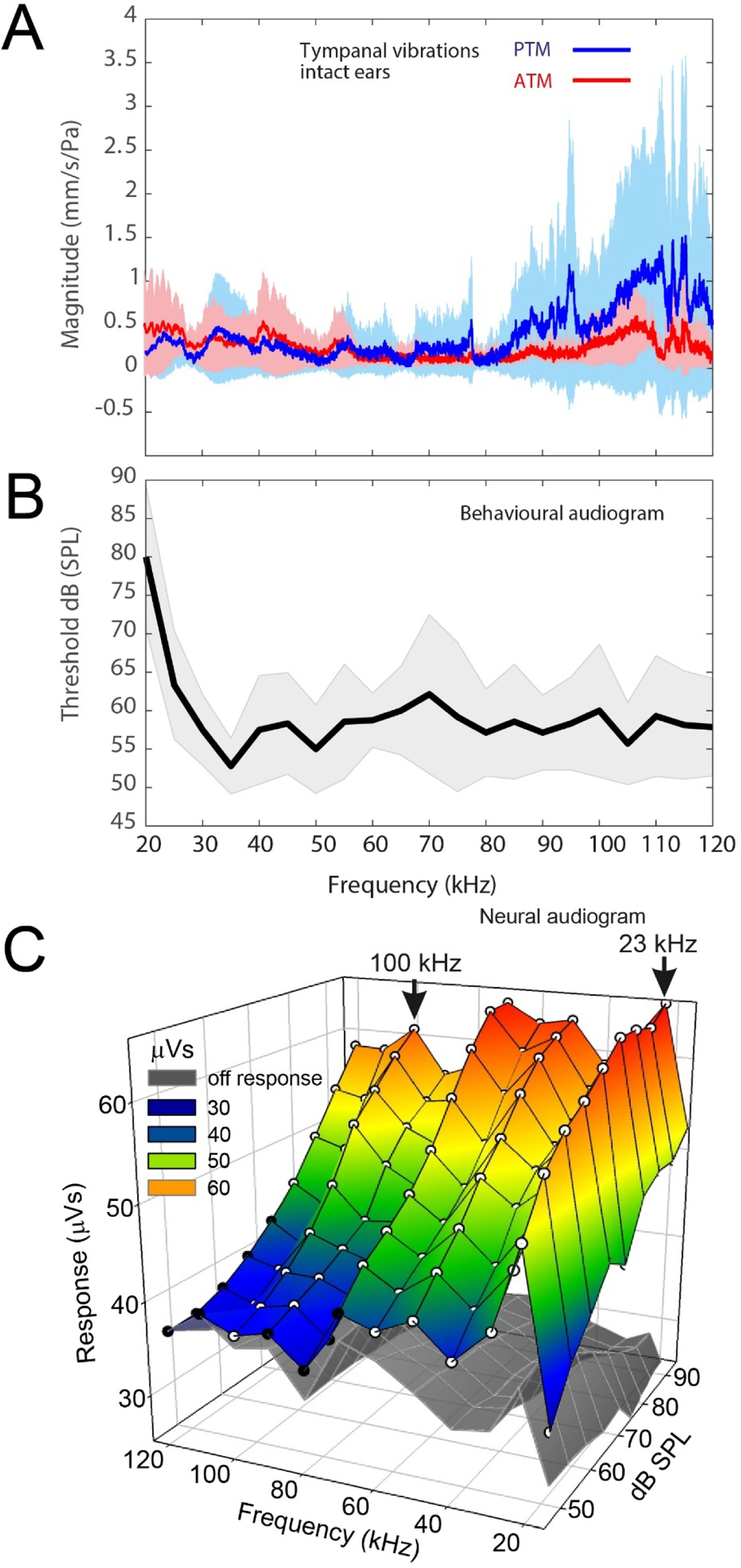
Tympanal tuning, behavioural and neural audiograms of *Copiphora gorgonensis*. ***A*.** Vibrational responses to broadband chirps (20 to 120 kHz) of real tympanal membranes (*n* = 7; 14 ears; four males and three females) of live *C. gorgonensis*. Maxima resonance peaks at 107.84 ± 3.74 kHz for the PTM and 111.13 ± 4.24 kHz for the ATM. Blue bar for PTM and red bar for ATM. ***B***. Black outline with grey shadow indicate the behavioural audiogram of ultrasound response in nine female *C. gorgonensis*. Black outline shows mean vector of SPL response at a particular frequency, shaded area represents the standard deviation across measured SPL. ***C***. Neural audiogram showing electrical activity in extracellular recordings of the auditory nerve during sound stimulation as sound pressure intensity (dB SPL) and sound frequency (kHz) are systematically altered. The coloured mesh shows the mean neuronal activity during the presentation of sound, and the grey mesh the mean neuronal activity during silent inter-pulse intervals. White symbols indicate a significant difference (*P* < 0.05) in paired t-tests comparing the sound-on and sound-off responses in each katydid tested; black symbols indicate a non-significant difference (*n* = 5). A strong neuronal response is apparent across a wide range of frequencies including to calling song (23 kHz) and to the high ultrasonic frequencies used by hunting bats (e.g., 100 kHz).

Behavioural audiograms of startle behaviour were obtained from nine tethered females walking on a treadmill. Audiograms were obtained with stimuli in the range 20 to 120 kHz. Audiograms show that the startle response of females decline sharply for stimuli between 20 kHz and 35 kHz, however, response remains essentially constant at higher frequencies over the entire tested frequency range (table S3; Fig. 5*B*).

### Neural audiograms

Auditory stimulation (the coloured mesh Fig. 5*C*) produced significantly greater responses in the neural audiogram recordings compared to neuronal activity during silent periods (grey mesh Fig. 5*C*) for most combinations of sound frequency and intensity (white symbol Fig. 5*C*). Only a small number of the stimuli failed to produce a significant difference in neuronal response, which occurred when frequency was high and sound pressures low (Fig. 5*C* black symbols). Within this region, responses to 100 kHz pulses were resolvable against background activity even for the quietest sound pressures (46 dB SPL).

Amplitudes of response increased almost linearly with sound pressure intensity across the frequency range. The largest responses were seen at the calling song frequency (23 kHz) across the sound pressure intensity range: at 46 dB the response was 51.4 ± 21.8 µVs, 35.6% greater than the activity during silence. At 70 dB SPL the response rose to 62.7 ± 15.8 µVs during stimulation, which was 86.1% higher than the equivalent off response, and rose further to 65.8 ± 14 µVs, (132.2% greater than the off response) at 94 dB SPL.

There was generally a gradual falling away of responsiveness as stimulus frequency increased above 23 kHz: at 70 dB SPL, the response to 40 kHz stimulation was 51.8 ± 12.2 µVs; the response to 60 kHz 49.1 ± 14.1 and to 80 kHz stimulation 44.5 ± 12.3 µVs, but measured responses to sound were still substantially above background activity. For example, at 100 kHz, the response of 47.5 ± 12.7 µVs was 43% greater than background activity. The weakest set of responses was to 120 kHz, which were not statistically distinguishable from the background until above 70 dB SPL, but nevertheless demonstrated that very high ultrasonic frequencies can be detected in *C. gorgonensis*.

## Discussion

To understand the function of cuticular pinnae of katydid ears, we conducted acoustic experiments on living specimens, combined with µ-CT to produce domains for numerical modelling using accurate ear geometries and to print 3D scaled ears for additional acoustic experiments. In all our experiments, pinnae had a significant effect on the sound reception at the tympana. Pinnae significantly enhanced cavity-induced pressure gains in live specimens at 60 kHz (the maximum frequency achieved with the experimental platform for living specimens). Further, the extent of the pinnal contribution to tympanal displacement amplitude depended on the incident angle of the sound source at tested frequencies ≤ 60 kHz. The tympana of *C. gorgonensis* naturally resonates at ca. 23 kHz, which shows high sensitivity to the dominant frequency of the male calling song (Celiker et al., 2020b; Montealegre-Z and Postles, 2010). This was also observed in our experimental results, irrespective of pinnal presence or absence. At ultrasonic frequencies (> 60 kHz), the pinnae-enclosed tympanal membrane of *C. gorgonensis* show strong mechanical vibrations induced by the resonances of the tympanal cavities (Fig. 5*A*). This suggests that tympanal pinnae enhance sound pressure and sensitivity to high frequencies. It was previously demonstrated that even minuscule tympanal displacements in *C*. *gorgonensis* create large displacement of the *crista acustica* (CA) (Montealegre-Z and Robert, 2015). Tympanal displacements are magnified in the CA as the effect of the lever action imposed by the vibration of the tympanum (Montealegre-Z et al., 2012). Comparable findings can be inferred from LDV experiments in the katydid *Mecopoda elongata,* in which the CA was exposed by microdissection (Hummel et al., 2014). Insect mechanosensory auditory neurons are capable of detecting exquisitely small mechanical displacements, down to 100 pm (Windmill et al., 2007), close to the theoretical limits of sensitivity (Bialek, 1987). Therefore, the sound pressure gain induced by the tympanal pinnae at ultrasonic frequencies (> 60 kHz; Fig. 3*A* and Fig. 5*A*) should produce sufficient tympanal displacement without EC amplification to induce a response in the auditory receptors. Electrophysiological recording of the auditory nerve from our experiments demonstrated a significant neural response to a broad range of frequencies (23–120 kHz) and sound pressures (46–94 dB SPL) (Fig. 5*C*; see *SI Appendix* Materials and Methods Section 1 for further details), demonstrating that *C*. *gorgonensis* can detect very high ultrasonic frequencies. Recordings from the T-cell in several katydids with pinnae (Faure and Hoy, 2000; ter Hofstede et al., 2010) and phaneropterines (ter Hofstede et al., 2010) without pinnae show a broad sensitivity in the range 5 to 85 kHz (see *Discussion*).

As a result of probe-speaker frequency range limitations and reflections from the specialised platform in our live time domain experiments, we were unable to test frequencies above 60 kHz. We compensated by performing numerical simulations on 3D computer geometries of experimental ears to predict pressure gains, 3D printing ear models to confirm resonance and pressure gain, and recording free field vibrating tympana in living experimental specimens. Our numerical simulations, considered in the frequency range 2-150 kHz, predicted sound pressure gains in the interval 50 to 150 kHz with resonant frequencies of 115 kHz and 121 kHz in the PTM and ATM cavities, respectively (Fig. 4*C*). Between 50 to 60 kHz, detectable pressure gains inside the cavity started to act along the external surface of the tympanal membrane. We did not compare the tympanal response between the experimental and numerical data directly, due to the simplifying assumption in the numerical models that the tympana are composed of homogeneous material. In reality, the tympana are composed of materially different layers where the external surface has more chitinous, sclerotized layers extending from the tympanal plate to the membrane, while the internal membrane is composed of elastic, tracheal derived material. To overcome this, we measured tympanal vibrations to extreme ultrasound in free field conditions to validate both the numerical and 3D print model results.

To test the influence of pinnal geometry alone on these ultrasonic gains, we printed 3D-scaled ears and scaled the sound stimuli to match the ear size. The mean resonance of the 3D printed models was found to be 101.47± 3.43 kHz for both the ATM and PTM (Fig. 4*E*). Differences between the numerical and 3D print models results can be attributed to the material properties (Young’s modulus of printing resin) and tympanal thickness incorporated for the tympana in the models. When we incorporated the material properties of the 3D ears into the mathematical models, the resonant frequencies dropped from 118 to 111 kHz (ATM 112.5 kHz, PTM 109.5 kHz; fig. S2) which is close to the high frequency resonance of the tympana when the pinnae are intact (107.84 ± 3.74 kHz for the PTM and 111.13 ± 4.24 kHz for the ATM; Fig. 5*A*). Without the pinnae, pressure gains were dramatically reduced in the simulations. Additionally, a slight resonance was found in both the numerical simulations and 3D print models caused by the incomplete ablation of the pinnal structures (Fig. 3*D* and *F*).

### Ultrasound guides in insects

Pinnae-covered tympanal ears are also found in some prominent moths (Notodontidae), with eardrums mechanically tuned to detect the high frequencies used by hunting bats (Windmill et al., 2006) which range from as low as 35 to above 135 kHz (Geipel et al., 2021). Cup-like pinnae from the metathorax are thought to enhance the reflection of sounds onto the tympanic membrane. With the pinnae ablated, these moths entirely lose the ability to localise sound at all frequencies (Fullard, 1984a). Therefore, katydids and certain moths have independently evolved pinnal structures for detecting ultrasounds.

High frequency singers of the katydid subfamily Pseudophyllinae generally have very small spiracles, long but narrow ECs, and tympana covered with various forms of cuticular pinnae (Heinrich et al., 1993). Pseudophyllines with song frequencies greater than 50 kHz have been shown to depend more on external than internal sound reception for communicating with conspecifics (Bailey et al., 1988; Bailey and Stephen, 1978; Mason et al., 1991; Stephen and Bailey, 1982). The dominant port for hearing relies on the tympanal slit to spiracle size area ratio where the larger input opening dictates principal auditory input in ultrasonic hearing rainforest pseudophyllines (Mason et al., 1991). In *C. gorgonensis*, the spiracle area is large, naturally open on average three times larger than the total area of the tympanal slits (1 mm^2^:0.3 mm^2^) which is inversely related to the general scale of pseudophyllines. Reflectance and power transmittance inside the tympanal pinnal cavities differ from the acoustic dynamics in the EC. Power transmittance of ultrasonic frequencies suffers significant attenuation as a result of high reflectance of sound waves along the narrowing EC in *C. gorgonensis* (Celiker et al., 2020a), limiting the EC response to frequencies <60 kHz.

By concentrating or funnelling ultrasonic sound into the tympanal cavity, the pinnae enhance ultrasonic reception of incidental sounds. The cavity induced pressure gains are the consequence of geometry of the tympanal slit in relation to the geometry and volume of the cavity (table S2). These imparted forces are magnified by the motion of the tympanum. The resonances afforded by the pinnal structures are evident as both the numerical and 3D print models do not include a vibrating tympanum. In *C*. *gorgonensis*, irrespective of incident sound pressure magnitude, the cavities provide a consistent pressure gain of at least 23 dB within the frequency range 100 to 120 kHz (Fig. 4*C*). This is in contrast to tympanate moths that depend on the incident sound intensity for mechanical tuning of high frequency bat calls (Fullard, 1984a, 1984b; Windmill et al., 2006) to produce gains up to 16 dB (Fullard, 1984a).

Though the slit openings to both cavities are perceptively indistinguishable from each other, the ATM slit opening is slightly larger than that of the PTM, but the PTM has a larger cavity volume. These minuscule discrepancies cause differences in pressure gains, resonances, and phase between vibrations of the ATM and PTM. In *C. gorgonensis*, the PTM pinna is approximately 13% wider than the ATM. This increases the micro-acoustical diffraction of ultrasonic frequencies entering the PTM cavity and potentially contributes to the broad turning of the PTM (Fig. 5*A*). Similar pinnal asymmetries were observed in the katydid *Oxyecous lesnei* but the ATM structure was much larger than the PTM which suggests that each tympanum is differentially tuned (for other examples see Fig. 1 (Sarria-S et al., 2017); fig. S3) (Bailey, 1993). Here, we showed that the tympanal cavities and their asymmetric openings act as Helmholtz resonators at frequencies of 94.13 ± 3.53 kHz for the ATM and 91.69 ± 3.93 kHz for the PTM in *C. gorgonensis*. Though the numerical simulations show an average peak of 118 kHz (ATM 121, PTM 115 kHz), the Helmholtz calculation assumes the slit opening is a circle and the cavity is a solid walled sphere. Pressure distribution maps from the numerical models suggest that at 110 kHz (see Fig. 4*A*) the resonances of the pinnae may function in a piston motion, whereby fluctuating air movements are present at the opening of each independent cavity: an observation characteristic of Helmholtz resonance. Quantitative imaging of scaled stimuli 9.63 kHz (110 kHz) using refracto-vibrometry in the area around the 3D printed ear (1:11.4) demonstrated such piston behaviour (*SI Appendix*, Supplementary Materials and Methods Section 2: Video 3). Sound pressure gains inside the 3D printed model tympanal cavities and resonances closely matched the simulated numerical models (Fig. 4*C* and *E*), showing an average sound pressure gain of 26.33 ± 4.06 dB for the ATM and 30.04 ± 1.34 dB for the PTM in the 3D printed ears at 101.47 kHz compared to 20.40 ± 2.82 dB for the ATM at 121 kHz and 22.92 ± 1.03 dB for the PTM at 115 kHz in the numerical models. When tympanal thickness and the material properties of the 3D print models were incorporated in the simulations, the average sound pressure for the ATM was 20.37 ± 3.52 dB at 112.5 kHz and 21.40 ± 2.57 dB at 109.5 kHz for the PTM (fig. S2).

The asymmetry of the tympanal cavities has an acoustic function. Our results show that pinnae cause intra-aural time differences and oscillation phase shifts at ultrasonic frequencies between vibrations in the ATM and PTM (Fig. 2*A*). This contributes to differences of intensity and arrival of sound that induce pressure gains of ultrasonic frequencies (Fig. 4*C* and Fig. 5*A*). Differential tympanal mechanical responses have also been found in the pinnae possessing paleotropical katydid *Onomarchus uninotatus* where the ATM exhibits tuning to the conspecific call and the PTM’s response is tuned to higher frequencies suggesting a possible use in predator detection (Rajaraman et al., 2013). The ATM and PTM of *C*. *gorgonensis* show differential responses to ultrasonic frequencies (Fig. 4*C* and 5*A*), but how these two signals are transduced in the same auditory sensilla for potential directional hearing remains unknown, as both membranes share the same CA. Antiphasic motion of ipsilateral tympana in *Mecopoda elongata* has been suggested to drive transduction by imparting mechanical stresses on the CA thereby stretching the sensory dendrites of mechanoreceptors (Vavakou et al., 2021). If true, our findings from living specimens show that pinnae cause consistently longer phase delays of ∼0.25 cycle (= 90°) between vibrations of the ATM and PTM at all tested frequencies than with the pinnae ablated. Not only do pinnal structures provide a long diffractive edge for sound waves, but the dorso-ventral asymmetry between the ATM and PTM pinnae increase phase differences that tune the ear (Vavakou et al., 2021). We argue that it is also possible for asymmetries within a single ear to function in the same way, including at frequencies emitted by hunting bats. However, our data do not support the idea of a single ear being directional at least in the range 20 to 60 kHz (which includes the katydid’s calling frequency at 23 kHz and first harmonic at 46 kHz; fig. S8). Considering that the tettigoniid ear is capable of resolving such small differences in time and intensity between the two ears (Bailey, 1993, 1990), four external inputs could resolve the direction of an approaching bat and evoke ultrasound-triggered defensive behaviour (Faure and Hoy, 2000). The behavioural and ecological relevance of potential directional hearing using a single ear constitutes an outstanding question and will be the subject of future studies.

In *C. gorgonensis*, the dual inputs of the spiracle and the four external ports (the pinnae) function as a sound pressure gain compensation system. As previously shown for *C*. *gorgonensis* (Celiker et al., 2020a), and in other species (Heinrich et al., 1993; Michelsen et al., 1994), the EC with its exponential horn geometry acts as a bandpass filter limited in providing pressure gains to high ultrasonic frequencies (> 60 kHz, for *C. gorgonensis*) (Lewis, 1974; Veitch et al., 2021) and is designed to enhance detection of the specific carrier frequency and associated harmonic contents (46 kHz for *C. gorgonensis*). The effect of the EC geometry in reducing sound velocity is as much as ∼20% or ca. 60 µs, coupled with the contralateral EC, collectively a total of eight inputs are possible with each causing a vibration with variable delays in four internal and four external tympanal surfaces to the same stimulus (Veitch et al., 2021). Though the reduction in velocity contributes exceptional binaural directional cues, the external port provides real-time sensitivity to exploit fading bat ultrasounds. Ear pinnae act as extreme ultrasonic guides that can resonate at the high frequency components at the start of the echolocation calls of bats. Hence the EC might be a less efficient method of bat detection as the angle of incidence accompanied by time delays could shorten reaction times and obfuscate the localisation of the predator. This suggests that the EC in katydids without ear pinnae should exhibit broadband responses (see last section of *discussion*).

### Bat detection by resonance

It has been shown that katydids form a key part of the diet of many insectivorous bat species in various regions of world (Arlettaz et al., 1993; Buchler and Childs, 1981; Davison and Zubaid, 1992; Fenton and Museum., 1975; LaVal and LaVal, 1980; Raghuram et al., 2015; Whitaker and Black, 1976; Zhang et al., 2005). However, such ecological interactions have been more intensively studied in the Neotropical regions. Gorgona Island, Colombia, is home to over 33 bat species with many still undescribed and underrepresented in wildlife inventories, including at least three substrate gleaning bats of the neotropical leaf-nosed bat family Phyllostomidae (Murillo G et al., 2014). Neotropical katydids have evolved sophisticated auditory features as strategies for survival against substrate gleaning bats (Belwood, 1990; Belwood and Morris, 1987; Nickle and Castner, 1995; ter Hofstede et al., 2017, 2010). The habitat of *C. gorgonensis* is in cluttered vegetation of the tropical forest understory (Montealegre-Z et al., 2014). In this environment, acoustic signals are heavily attenuated which leads to significant transmission loss (Rheinlaender and Römer, 1986; Wiley and Richards, 1978). However, insects have evolved a variety of sophisticated receivers to perform call discrimination in these acoustically challenging environments (Römer, 1993). Acoustic adaptations by katydids to evade bat predation include the use of narrow bandwidths, high carrier frequencies, and sporadic calling in order to diminish signal proliferation in the environment (Belwood and Morris, 1987; Morris et al., 1994; Morris and Beier, 1982; Rentz, 1975), and ergo evade eavesdropping by bats (Heller, 1995). Certain adaptations are a trade-off as the katydid becomes more conspicuous and vulnerable to other predators as the communication method changes. For example, katydids that perform vibrotaxis, even tremulations can likely attract spiders, scorpions (Robinson and Hall, 2002) and primates, as well as bats (Geipel et al., 2020). Likewise, bats foraging at this level also face similar acoustic shortcomings, affecting their echolocation abilities (Page and Bernal, 2020). Thus, several phyllostomid substrate gleaning bats are very well adapted to listen to prey-produced sounds like rustling noises or mating calls of, e.g., male katydids (Belwood and Morris, 1987; Falk et al., 2015; Geipel et al., 2021). At least one common gleaning bat species, *Micronycteris microtis* (Phyllostomidae), uses a sophisticated echolocation strategy to detect katydids concealed in vegetation (Geipel et al., 2019, 2013). Despite their passive acoustic defences, calling from restrictive locations and being equipped with very large mandibles and sharp fastigia, katydids like *C. gorgonensis* are predated by phyllostomid bats (ter Hofstede et al., 2017).

Here we argue that the pinnal structures of the external tympanal ports of katydid ears act as sound guides providing acute ultrasonic hearing allowing the detection of echolocation calls of hunting bats and thus an additional sensory based defence in the predatory-prey arms race. The presented numerical and experimental evidence suggests that the greatest ultrasonic gain of the pinnae is at frequencies matching those falling into the frequency range of the echolocation calls of native, insect gleaning bat species (Fig. 6). As neotropical gleaning bats approach their target, they emit short, broadband, multi-harmonic sweeps, demodulate the frequency from higher frequencies above 135 kHz to as low 35 kHz (Geipel et al., 2021; Yoh et al., 2020). In terms of predator detection, a katydid like *C. gorgonensis* has an excellent chance of detecting the calls of a hunting bat even at the start of the sweep. Responses to these high frequencies are supported by LDV recordings of tympanal motion in intact ears of free-field specimens, and audiograms that show a broad mechanical, behavioural, and neural response to ultrasonic frequencies (table S3, Fig. 5*A-C*). These broad responses to ultrasound are common in several pinnal bearing katydids subfamilies of Tettigoniidae (Deily and Schul, 2006; Schul and Patterson, 2003), as shown in the Tettigoniinae with *Tettigonia viridissima* (Rheinlaender and Römer, 1986), in Pseudophyllinae with *Acanhodis curvidens* (ter Hofstede et al., 2010), *Balboa tibialis* (ter Hofstede et al., 2010), *Docidocercus gigliotosi* (ter Hofstede et al., 2010) *Ischnomela gracilis* (ter Hofstede et al., 2010), and in Conocephalinae for example *Bucrates capitalus* (ter Hofstede et al., 2010), *Neoconocephalus ensiger* (Faure and Hoy, 2000) and *Neoconocephalis affinis* (ter Hofstede et al., 2010). A gain of 16 to 20 dB at the start of the bat call provides essential awareness time (≤ 0.86 ms in terms of duration of the complete sweep (Geipel et al., 2021)) to *C*. *gorgonensis* as a result of the tympanal pinnae. Other katydid species living in sympatry with *C*. *gorgonensis* like *Supersonus aequoreus* (the most ultrasonic katydid found in nature to date (Sarria-S et al., 2014)), *Ischnomela gracilis*, and *Eubliastes aethiops* exhibit similar cavity induced pressure gains in the range of phyllostomid echolocation calls (fig. S4).

**Figure 6.**
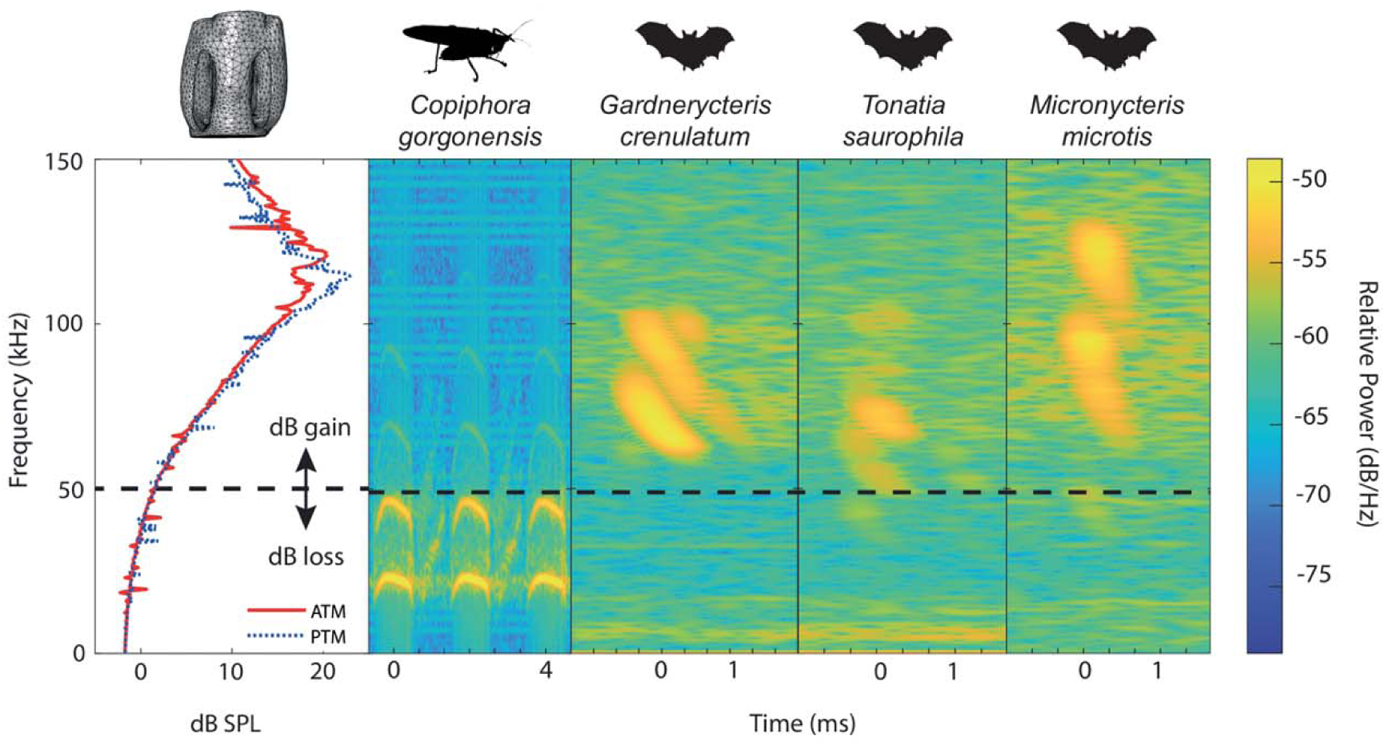
Ecological relevance of pinnae in *Copiphora gorgonensis*. Sound pressure level gains (left) induced by the pinnae are present only at frequencies above c.a. 50 kHz, covering the range of echolocation frequencies of three native insectivorous gleaning bat species. The conspecific call of *C. gorgonensis* (dominant frequency and harmonics) on the other hand (dB_peak_ at 23 kHz), is not enhanced by the presence of the pinnae (dB loss). Dotted line indicates the frequency at which gain = 0 dB. Spectrogram parameters: FFT size 512, Hamming window, 50% overlap; frequency resolution: 512 Hz, temporal resolution: 0.078 ms.

The pressure – time difference receiver of *C*. *gorgonensis* is a unique system that can capture different ranges of frequencies between the multiple entry ports that can obviate the limitations of each but is also capable of compensating for limitations in auditory orientation (Veitch et al., 2021). For katydids, incident sounds from elevation are difficult to perceive (Römer, 2020). The ability of the external port to be positioned and rotated in accordance with the movement of the foretibial knee and foretibial leg joints permits for the ear to be more vertically oriented. The µ-CT imaging presented here of the tympanal cavities supports the theory for vertical orientation as the sub-slit cavity is asymmetrically recessed to the distal end, which is likely to enhance mechanical responses to vertical stimuli (Fig. 1*D*). For ultrasonic reception, a total of four asymmetrical external ports (left and right ATMs and PTMs) may be behaviourally articulated in a manner to enhance the detection of bats calling from elevated positions towards the katydids, and the physical separation between the external ports of each ear yields sufficient binaural cues.

### Ideas and Speculation: Katydid ear pinnae and the fossil record

The presence of ear pinnae in katydids is unknown in the fossil record. Katydid ancestors (e.g., Haglidae and Prophalangopsidae from upper Jurassic (Plotnick and Smith, 2012)) and early katydids (Tettigoniidae) from the middle Paleogene (Eocene, ∼55 mya (Greenwalt and Rust, 2014; Rust et al., 1999)) all show naked tympana (without pinnae). The earliest echolocating bats are from the Eocene (Teeling et al., 2005), therefore katydids with tympanal pinnae may have initially evolved such sophisticated hearing devices to survive nocturnal predators while they sing under the cover of darkness. Although katydid ear pinnae have never been mapped in the most recent molecular phylogenies (Mugleston et al., 2018, 2013), we observe a potential unique origin of ear pinnae in the family Tettigoniidae, with multiple losses or retrogressions in modern species, including the large subfamily Phaneropterinae (predominantly known to have naked tympana). Comparative analyses using large phylogenies are needed to solve this working hypothesis. Analogous ear pinnae adaptations are observed in some Eneopterinae crickets (tribe Lebinthini) (Schneider et al., 2017), which differ from field crickets in their use of high frequencies for specific communication (12 to 28 kHz). These crickets emerged also in the Eocene (Vicente et al., 2017) and while their ancestors exhibit only one functional tympanum (PTM), the extant forms show two functional asymmetric tympana, with the ATM covered by pinnae (Schneider et al., 2017). Such adaptation suggests a new paradigm of the dual role of the ears, in detecting conspecific and bat echolocation calls.

Several katydid species (e.g., Phaneropterinae, Mecopodiae) exhibit naked tympana (fig. S1*B*). While little is known about the ecology of many species, katydids have developed diverse hearing structure morphologies to respond to predation pressure (Hoy, 1992). Ears evolve very rapidly (Hoy, 1992; Song et al., 2020) and it would not be surprising that, without pinnal structures, some nocturnal Phaneropterinae evolved sophisticated ECs with exceptional broadband response (i.e., broader than that of *C*. *gorgonensis*) (Heinrich et al., 1993; Hoffmann and Jatho, 1995; Michelsen et al., 1994; ter Hofstede et al., 2010). A study from Barro Colorado Island, Panamá, reported 31 Phaneropterinae katydids, of which about 42% use calling songs in the low ultrasonic range while 74% showing a spectral bandwidth of >10 kHz (ter Hofstede et al., 2020). It might also be possible that some Phaneropterinae have a unique ear function via the EC which is capable of detecting conspecific calls as well as bats. This configuration of a single ear function is also potentially exhibited by *Supersonus aequoreous* (fig. S3) which show atrophied EC, and their outer ear components (tympana and pinnae) specialised in identifying the direction of their own calls while at the same time detecting bats. Other adaptations of katydids without tympanal pinnae for bat detection might involve activity during the daytime (Hoy, 1992; ter Hofstede et al., 2020), or dwell in dense vegetation that challenges hunting bats (Lang and Römer, 2008). For example, *Phlugis poecila* and *Speculophlugis hishquten* species are diurnal visual predators that need day light to hunt (Woodrow et al., 2019). Owing to their transparent camouflage, males are able to sing below Araceae leaves during the day and avoid visual detection by diurnal avian predators. Other katydids like *Conocephalus* sp. with tympanal pinnae are also active during the daytime, although a few species are nocturnal or crepuscular, but a majority dwell in dense grass vegetation. Their calling songs are of unusual broadband energy, in many species expanding above 60 kHz (Fullard et al., 1989). In this case, the retention of the pinnal condition might be associated with specific directional hearing like acoustic ranging (Harness and Campbell, 2021) in such dense grass environments (Rheinlaender and Römer, 1986; Romer and Lewald, 1992).

While the diversity of form and function of pinnae in katydids requires a deeper comparative analysis, the presented findings suggest that in the assessed species, pinnae act as ultrasonic resonators for the early detection of echolocating bats. As a working hypothesis, we propose that the ear pinnae have a unique origin across the ca. 8000 living species of Tettigoniidae (Cigliano et al., 2021) in response to the emergence of bats during the early Eocene, and that it was subsequently lost or modified several times.

## Materials and Methods

### Specimens

*Copiphora gorgonensis* (Tettigoniidae: Copiphorini) is endemic to Gorgona National Natural Park, Colombia (02°58′03″ N 78°10′49″W). The original generation of the species were imported to the UK under the research permit granted by the Colombian Authority (DTS0-G-090 14/08/2014) in 2015. The specimens were ninth generation, captive bred colonies and maintained at 25°C, 70% RH, light: day 11 h: 23 h. They were fed *ad libitum* diet of bee pollen (Sevenhills, Wakefield, UK), fresh apple, dog food (Pedigree Schmackos, UK) and had access to water. Live experiments were conducted on seven adults of *C. gorgonensis* from our laboratory breeding colonies at the University of Lincoln (Lincoln, UK). Following experimentation, these specimens plus an additional four females already stored in ethanol were µ-CT scanned for finite element modelling; totalling 17 ears (10 female, 7 male). Live specimens were subsequently preserved in 100%ethanol-filled jars and stored in a freezer at −22°C at the University of Lincoln (Lincoln, UK).

### Simultaneous recordings of tympanal vibrations using laser Doppler Vibrometry

Insects were chemically anesthetized using triethylamine-based agent FlyNap (Carolina Biological Supply, USA) for 15 min prior to the mounting process, and remained awake throughout the duration of the experiment. The animals were dorsally mounted using a specialized platform to isolate the external and internal sound inputs and also mimic their natural stance (fig. S5). A rosin-beeswax mix was used to fix the pronotum, and the mid and hind legs, to the mount. This specialized platform (Jonsson et al., 2016) consists of two Perspex panels (1.61 mm thick) that are joined by latex and suspended in the air by a 12 x 12 mm metal frame attached to a micromanipulator (World Precision Instruments, Inc., USA) (see (Montealegre-Z et al., 2012)). At the Perspex junction, the forelegs of the insect were extended through arm holes cut in the Perspex and attached on a rubber block with metal clasps. A metal clasp was placed on each foretibia and forefemur (total of 4) to arrest foreleg motion. The arm holes and frame borders were sealed with latex to deny sound propagation to the spiracle.

The laser Doppler vibrometry system consisted of a the OFV-2520 Dual Channel Vibrometer - range velocity controller for operating two single point laser sensor heads, (OFV-534, Polytec, Germany) each with VIB-A–534 CAP camera video feed and laser filters. Each sensor head was mounted on a two-axis pivoting stage (XYZ, Thorlabs Inc., USA) anchored to an articulating platform (AP180, Thorlabs Inc., USA) and manually focused at 10.5 cm above a vibration isolation table (Pneumatic Vibration Isolation Table with a B120150B - Nexus Breadboard, 1200 mm x 1500 mm x 110 mm, M6 x 1.0 Mounting Holes, Thorlabs Inc., USA) supported by an anti-vibration frame (PFA52507 - 800 mm Active Isolation Frame 900 mm x 1200 mm, Thorlabs Inc., USA) in an anechoically isolated chamber (AC Acoustics, Series 120a, internal dimensions of 2.8 m x 2.7 m x 2 71 m). The sensor heads were outfitted with magnification microscopic lenses (Mitutoyo M Plan 10x objective for Polytec PSV-500 single laser head OFV 534, Japan) and positioned about 35 to 40 mm away from the insect foreleg at 45° angles towards the Perspex surface (fig. S5). The narrow entrance to the tympanal cavities restricted the use of LDV to measure tympanal responses across the entire tympanal membrane. Therefore, the placement of the sensor heads was limited to positions where the sensor heads were perpendicular to the tympanic membrane of interest. The sensor speeds were maintained at 0.005 (m/s)/V and recorded using an OFV-2520 internal data acquisition board (PCI-4451; National Instruments, USA).

Tympanal vibrations were induced by a four-cycle sinusoidal wave at 23, 40 and 60 kHz. The closed-field configuration of the loudspeaker restricted the delivery of high ultrasonic stimuli to 60 kHz. A rotating automated stage (PRM1Z8 rotation mount, Thorlabs Inc., USA) with a KDC101 K-Cube™ DC Servo Motor Controller (Thorlabs Inc., USA) positioned a multi-field magnetic loudspeaker (MF1, Tucker Davis, USA) with a parabolic nozzle (see Supplementary Materials from (Veitch et al., 2021)) and plastic probe tip (3.5 cm L x internal diameter 1.8 mm W) about 3.5 mm away from the mounted insect and 10.2 cm above the breadboard table. The speaker was moved across a 12 cm semi-circle radius in 1° steps (0.56 mm). The probe tip was positioned at “point zero” and 20 single shot recordings at 1°intervals, totalling 10° at either side (fig. S6). A high quality 500 band pass filter was applied at 10 to 30 kHz for the 23 kHz recordings, 30 to 50 kHz for the 40 kHz recordings, and 50 to 70 kHz for the 60 kHz recordings. All acoustic signals were generated by a waveform generator (SDG 1020, Siglent, China), synchronized with the LDV, amplified (ZB1PS, Tucker Davis, USA) and measured by a 1/8 (3.2 mm) omnidirectional microphone (Type 4138, Brϋel & Kjaer, Nærum Denmark) located about 3 mm from tympanum. The microphone, with built in preamplifier (B&K 2670, Brüel & Kjær, Nærum, Denmark), was calibrated using a sound-level calibrator (Type 4237, Brϋel & Kjaer, Nærum, Denmark) and set to 316 mV/Pa output via a conditioning amplifier (Nexus 2690-OS1, Brüel & Kjær, Nærum, Denmark). A reference measurement was performed by placing the microphone 3 mm from the probe tip to the loudspeaker before each experiment. Using a micro-manipulator, the microphone was positioned approximately 3 to 3.5 mm from the ear to monitor the acoustic isolation of the platform.

### Experimental procedures

The sensor heads were manually focused on the external tympanal surface using the 2-axis pivoting stage and manual wheel with the aid of the sensor head camera output displayed on an LED screen. For the time measurements, the point zero was found for each leg and for each test frequency. The point zero was the point where the displacements from the anterior tympanic membrane (ATM) and posterior tympanic membrane (PTM) matched the oscillation phase of the generated 4-cycle sinusoidal waves. This ensured that the vibrations of the tympanic membranes were synchronous relative to the speaker position. Displacement amplitudes from the same cycle order number were measured from each sensor head reference, and approximately 252 data points were measured per ear.

After recording the vibrations for both ears of the tested individual, the cuticular pinnae were carefully excised using a razor blade (taking care not to damage the tympanal organs or the fine layer of tissue ventrally connected to the tympanic membranes). The measurements were repeated for each ear following the same protocol.

Time and displacement measurements were analysed by identifying the second oscillation of the 4-cycle tone generated waves in each software window (PSV 9.4 Presentation software, Polytec, Germany). Phase calculations were obtained using the equation φ° = 360° *x f x Δt* where *f* is frequency (kHz) and Δ*t* (ms) the difference in arrival times between the ATM and PTM.

### Morphological studies of the ear

To produce 3D data for modelling, 17 ears of *C. gorgonensis* were scanned using a SkyScan 1172 X-ray µ-CT scanner (Bruker Corporation, Billerica, MA, USA) with a resolution between 1.3 and 2.9 µm (55 kV source voltage, 180 µA source current, 300 ms exposure and 0.1° rotation steps). As experimental procedures required removal of the cuticular pinnae, eight additional specimens with intact pinnae were scanned. The µ-CT projection images were reconstructed with NRecon (v.1.6.9.18, Bruker Corporation, Billerica, MA, USA) to produce a series of orthogonal slices. The 3D segmentation of the ear, measurements of the ear cross section and width, and volumetric measurements of the cavities formed by the pinnae were performed with the software Amira-Aviso 6.7 (Thermo Fisher Scientific, Waltham, Massachusetts, USA). µ-CT stereolithography files (STL) were generated for numerical modelling using established protocols (Jonsson et al., 2016; Veitch et al., 2021) and to 3D print ear models.

For 2D measurements of the cavity slit area, pinnal protrusion, and the distance between the pinnal cavities, an Alicona InfiniteFocus microscope (G5, Bruker Alicona Imaging, Graz, Austria) at x5 objective magnification was used to capture images of collection specimens with intact pinnae, with a resolution of about 100 nm (*n* = 8 ears).

### Bat and insect call recordings

The echolocation calls of phyllostomid bats (Chiroptera: Phyllostomidae) native to Gorgona Island (*Gardnerycteris crenulatum*, *Tonatia saurophila* and *Micronycteris microtis*) were recorded in a small indoor flight cage (1.4 x 1.0 x 0.8 m) in which they were allowed to fly, via an ultrasound condenser microphone (2-200 kHz frequency range, ±3 dB frequency response between 25-140 kHz; CM16, CMPA preamplifier unit, Avisoft Bioacoustics, Glienicke, Germany) and real time ultrasound acquisition board (6 dB gain, 500 kHz sampling rate, 16 bit resolution; UltraSoundGate 116Hm, Avisoft Bioacoustics, Glienicke, Germany) connected to a laptop (Think Pad X220, Lenovo, Beijing, China), with a corresponding recording software (Avisoft RECORDER USGH, Avisoft Bioacoustics, Glienicke, Germany). These are the most common insectivorous gleaning species in the habitat of *C. gorgonensis*. Single calls of each species are presented in Fig. 6 and fig. S7.

*SI Appendix*, Section 1 (see fig. S8) has details of male *C*. *gorgonensis* calling song recording.

### Acoustics measurements of synthetic 3D-printed scaled ear models

For time domain measurements, 3D models of the ears were placed on a micromanipulator arm with blu-tac (Bostik Ltd, Stafford, UK) and positioned frontally 30 cm from a MF1 loudspeaker at the same elevation. A 25 mm tipped B&K Type 4182 probe microphone (Brüel & Kjær, Nærum, Denmark) with a 1 × 25 mm (0.99″) probe tube length and 1.24 mm (0.05″) interior diameter, calibrated using a B&K Type 4237 sound pressure calibrator was placed ventral to the ear. The ear moved on the microphone using an electronic micromanipulator (TR10/MP-245, Sutter Instrument, Novato, California, USA), to a position 1 cm from the back of the cavity. Stimuli delivered were individually scaled to match the wavelength of a real-size ear (e.g., for a 1:10 scale printed model, the frequency delivered to simulate 120 kHz would be 120/10 = 12 kHz) to account for variation in printed model scaling. 3D printed models were scaled 1:11.43 (male 1:11.33; female 1:11.53) with the corresponding average scaled stimuli of 2.01 kHz for 23 kHz, 3.50 kHz for 40 kHz, 5.25 kHz for 60 kHz, and 9.63 kHz for 110 kHz. Four cycle pure tones were produced using the aforementioned function generator, and the amplitude set to deliver 1 Pa to the microphone at each frequency. Received signals were amplified using a B&K 1708 conditioning amplifier (Brüel & Kjær, Nærum, Denmark), and acquired using a PSV-500 internal data acquisition board at a sampling frequency of 512 kHz. The microphone remained stationary during the experiments, nor was its direct path to the speaker obstructed. Instead, the microphone entered the ear via a drilled hole, allowing the pinnae to surround the tip of the microphone. Thus, the reported sound pressure gains result solely from the cavities of the 3D model, and not the motion of the microphone. When the microphone was positioned inside the cavities, the gap between the drilled hole and microphone probe was sealed with blu-tac (Bostik Ltd, Stafford, UK) to mimic the real cavity and avoid acoustic leaking (refer to *SI Appendix*, Supplementary Materials and Methods Section 2: Video 1; Video 2).

To calculate the frequency that produced the best gain, the MF1 loudspeaker was replaced with a RAAL 140-15D Flatfoil loudspeaker (RAAL, Serbia), with a different amplifier (A-400, Pioneer, Kawasaki, Japan). This speaker was able to deliver a broadband stimulus of periodic chirps, generated within Polytec 9.4 software, with a simulated frequency range of 2 to 150 kHz. Recording in the frequency domain, at a sampling frequency of 512 kHz, the amplitude of the broadband stimulus was mathematically corrected within the software to deliver 60 dB at all frequencies. The reference frequency spectrum with no ear present could be subtracted from the frequency spectrum reported within the cavities to calculate frequency-specific gain and thus cavity resonance. Gain was calculated by subtracting the probe microphone sound pressure (dB) measured 1 cm outside of the cavity from inside the tympanal cavity measurements (Fig. 3*B* and 3*C*; see also *SI Appendix*, Supplementary Materials and Methods Section 2: Video 1). For comparative purposes, the ears of the following sympatric and pinnae bearing katydid species from Gorgona Island were also 3D printed and subjected to experiments according to the aforementioned protocol: *Ischnomela gracilis*, *Supersonus aequoreus* and *Eubliastes aethiops* (see fig. S3).

Frequency domain recordings of the cavity resonance, and time domain recordings of pure tone gains were then exported as .txt files for analysis. Methods for printing 3D ear models are provided in the Supplementary Materials and Methods Section 1 of the *SI Appendix*.

### Mathematical models and numerical simulations

The mathematical models have been constructed as a scattering acoustic – structure interaction problem and simulate the acoustic response of the tympanal cavities to an incident plane acoustic wave in an air domain. Hence, the 3D model considers the interaction of the sound wave with the ear, for which realistic material properties have been incorporated. The air acoustic domain is truncated as a sphere with a 3 mm radius that is centered around the ear (fig. S9). Two different geometries of the ears were taken as part of the mathematical model domain: pinnae intact and pinnae removed (fig. S10).

The models were considered both in the frequency and the time domains, and were solved using the acoustic-shell interaction module of the software Comsol Multiphysics, v5.6 (Comsol, n.d.). For the frequency domain models, the incident wave was taken to be a chirp with an amplitude of 1 Pa and frequency 2 to 150 kHz, directed at “point zero” as defined in the in the section *vibrational measurements*. For the time domain models, three different incident waves were used, with amplitudes 1 Pa and frequencies 23, 40, 60 kHz. The direction of the waves was taken as −10°, −5°, 0°, 5° and 10° on a fixed plane perpendicular to the ear, with 0° corresponding to “point zero”. The details of the solved system of equations can be found in the *SI Appendix* Supplementary Materials and Methods Section 1.

The numerical solution to the problem was obtained using the finite element method for the spatial variables in both the time and frequency domain simulations. For forming the finite-element mesh, the maximum diameter used for the tetrahedral elements in the sphere was 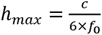, where *c* = 343 m/s and *f*_0_ = 150 kHz (figs. S11 and S12). Hence, even at the largest frequency considered, there were six tetrahedral elements per wavelength. Quadratic Lagrange elements were applied for the solution.

For the time domain solution, the time variable was solved for using the Generalized alpha method, with a constant time step of 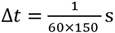 so that the Courant-Friedrichs-Lewy (CFL) condition (Courant et al., 1967), defined as 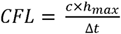 was 0.1, which gives a reliable approximation of the solution.

### Behavioral and tympanal response to broadband stimulation

Behavioral audiograms were measured from nine tethered female (*n* = 9) *C*. *gorgonensis* to test behavioral response thresholds to controlled auditory stimuli (20 to 120 kHz). Methods are provided in the *SI Appendix* Supplementary Materials and Methods Section 1.

For the tympanal tuning measurements, we exposed seven specimens(4 males, 3 females) to free field broadband (periodic chirp 20 to 120 kHz) stimulation presented by an ipsilaterally positioned SS-TW100ED Super-Tweeter (Sony, Tokyo, Japan) with a 20 kHz built-in high-pass filter using an Avisoft Bioacoustics Ultrasonics Power Amplifier (Avisoft Bioacoustics, Glienicke/Nordbahn, Germany). A rosin-beeswax mix was used to fix the pronotum, and the mid and hind legs, to the mount (see (Montealegre-Z et al., 2012)) after the insects were chemically anesthetized using FlyNap. Insects were then elevated to the same level as the LDV and positioned 15 cm from the loudspeaker. A 1/8” B&K Type 4138 microphone was placed about 3 mm in front of the ear of interest and recorded the stimulus. Mechanical responses were acquired using a PSV-500 internal data acquisition board at a sampling frequency of 512 kHz. The amplitude was corrected to maintain 60 dB SPL at all frequencies. Data was collected as magnitude (velocity/sound pressure).

### Neural audiograms

Extracellular recordings of whole auditory nerve activity in response to sound were obtained from five animals, using a suction electrode positioned on the nerve in the distal femur. Recordings were digitized using a Micro1401 mk II (Cambridge Electronic Design (CED), Cambridge, UK) for observation and storage for later analysis on a PC using Spike2 (CED) software. Stimuli consisted of 10 repeats of 500 ms sound pulses followed by 500 ms silent periods. Frequency (11 pure tones ranging from 23–120 kHz) and SPL (9 levels ranging from 46–94 dB in 6 dB increments) were systematically altered for a total of 99 combinations. Recordings for each train of 10 stimuli were root-mean-square transformed (time constant 0.66 ms) to convert the neuronal traces into positive displacements from zero and averaged. The mean areas of response (in microvolt s, µVs) during the presentation of each different sound stimulus was compared to the mean response during the subsequent silent period in each animal (*n* = 5) using paired *t*-tests. Detailed methods for neural audiograms are provided in the *SI Appendix* Supplementary Materials and Methods Section 1.

### Statistical analyses

Using empirical data we tested the effect of cuticular pinnae on tympanal responses [in displacement amplitude (natural log transformed) and arrival time] to incident sound, we fitted linear mixed models (LMM) with angle (−10° to 10°, quadratic polynomial continuous variable) as a covariate and presence of pinnae (y/n), frequency (23, 40 and 60 kHz, categorical variable), tympanum (ATM or PTM) as fixed factors. We included the interactions between angle and pinnae presence and between pinnae presence and frequency. To model the curvature in the response surface of the pinnal enclosed tympanum, angle was fitted as a quadratic polynomial with 0° at point zero. The interaction of angle and pinnae was fitted as such to show the restriction of pinnal structures in both time and displacement to the response surface. To account for repeated measures of the same specimen, we nested leg (left or right) within individual specimens as a random factor. We carried out post hoc tests between pinnae (y/n) at each frequency using estimated marginal means from the package *emmeans* (Lenth and Lenth, 2018).

Using the same initial LMM model, we tested how sound pressure estimated from numerical models was related to angle (−10° to 10°, polynomial continuous variable), presence of pinnae (y/n), frequency (23, 40 and 60 kHz, categorical variable), tympanum (ATM or PTM) as fixed factors. Again, we include the interactions between angle and pinnae and between pinnae and frequency. We finally tested sound pressure based on 3D models with the presence of pinnae (y/n), frequency (23, 40 and 60 kHz, categorical variable), tympanum (ATM or PTM) as fixed factors, and with the inclusion of the interaction between pinnae and frequency. For both numerical and 3D models, we carried out post hoc tests between pinnae (y/n) at each frequency using estimated marginal means from the package *emmeans*.

Statistical tests and graphs were performed in R 4.0.0 (Team, 2021) and all LMMs were run using the package lmerTest (Kuznetsova et al., 2017).

### Data availability

Experimental data (LDV recordings), numerical simulations, Comsol model files, and µ-CT stereolithography files (in .stl format) are available in Dryad (https://doi.org/10.5061/dryad.k0p2ngf8x; deposited 9 September 2021).

## Supporting information

Supplemental Index

## Acknowledgments

This research is part of the project “The Insect Cochlea” funded by the European Research Council, Grant ERCCoG-2017-773067 to FM-Z; and was also funded by the Natural Environment Research Council (NERC), grant DEB-1937815 to FM-Z. We thank the Orthopterists’ Society for aiding in funding the µ-CT work of CW, for which some data has been used in this study, and to the University of Lincoln’s School of Life Sciences for CW’s PhD studentship. The authors gratefully acknowledge Dr. Berthold Hedwig for sharing laboratory space, equipment and valuable advice.

## Supplemental Figures and Tables

**Figure S1.**
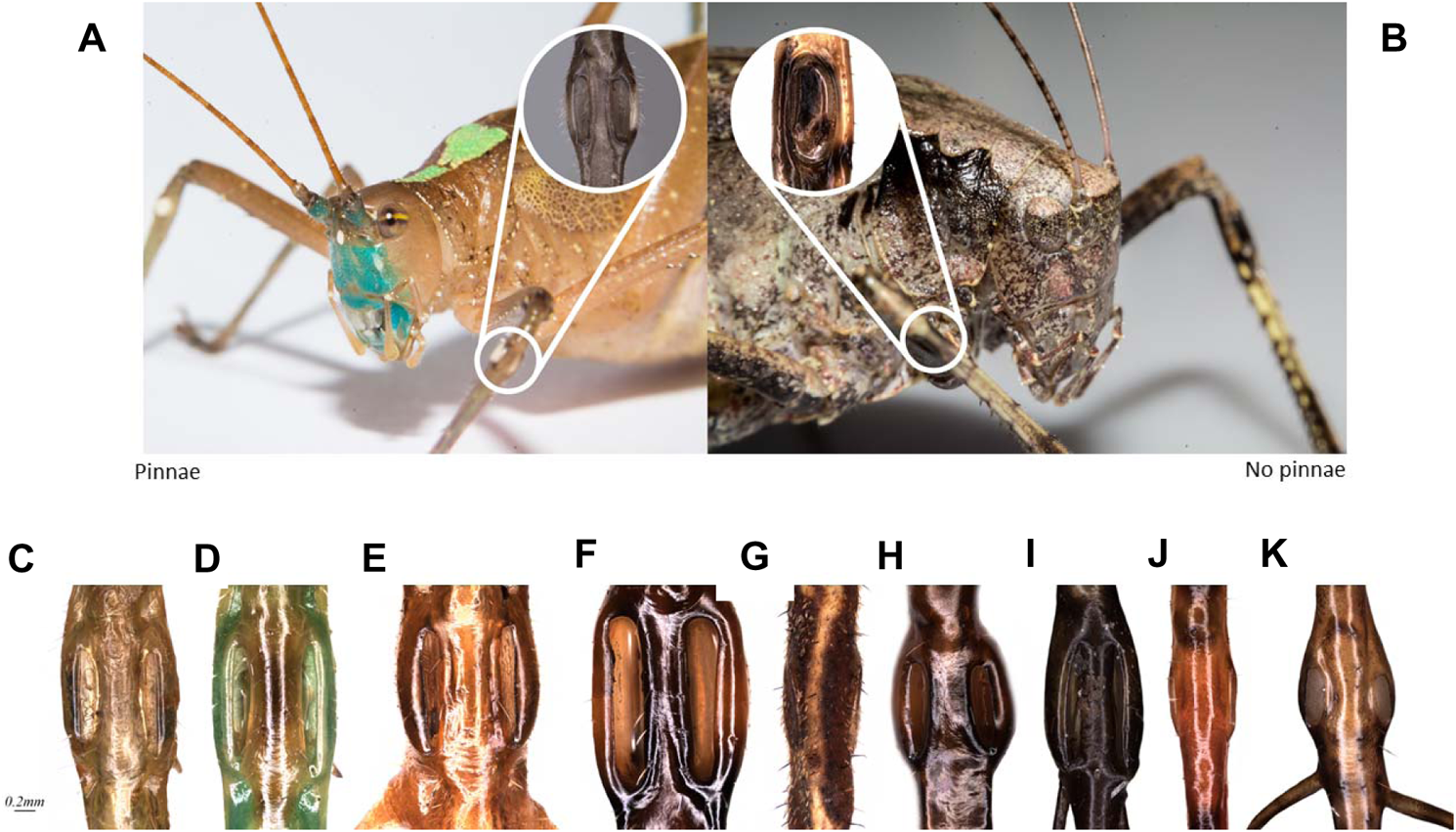
Cuticular pinnae of various neotropical katydids. **A**. Comparison between the pinnal ear of *Docidocercus sagittatus* and the naked ear of *Mecopoda elongata*. **C-K**. Pinnae bearing katydids: (**C**) *Copiphora gorgonensis*, (**D**) *Copiphora vigorosa*, (**E**) *Panacanthus pallicornis*, (**F**) *Cocconotus zebra*, (**G**) *Typophyllum spurioculis*, (**H**) *Acanthacara acuta*, (**I**) *Arachnoscelis arachnoides*, (**J**) *Paranelytra* sp. (undescribed sp.), (**K**) *Supersonus aequoreus*.

**Figure S2.**
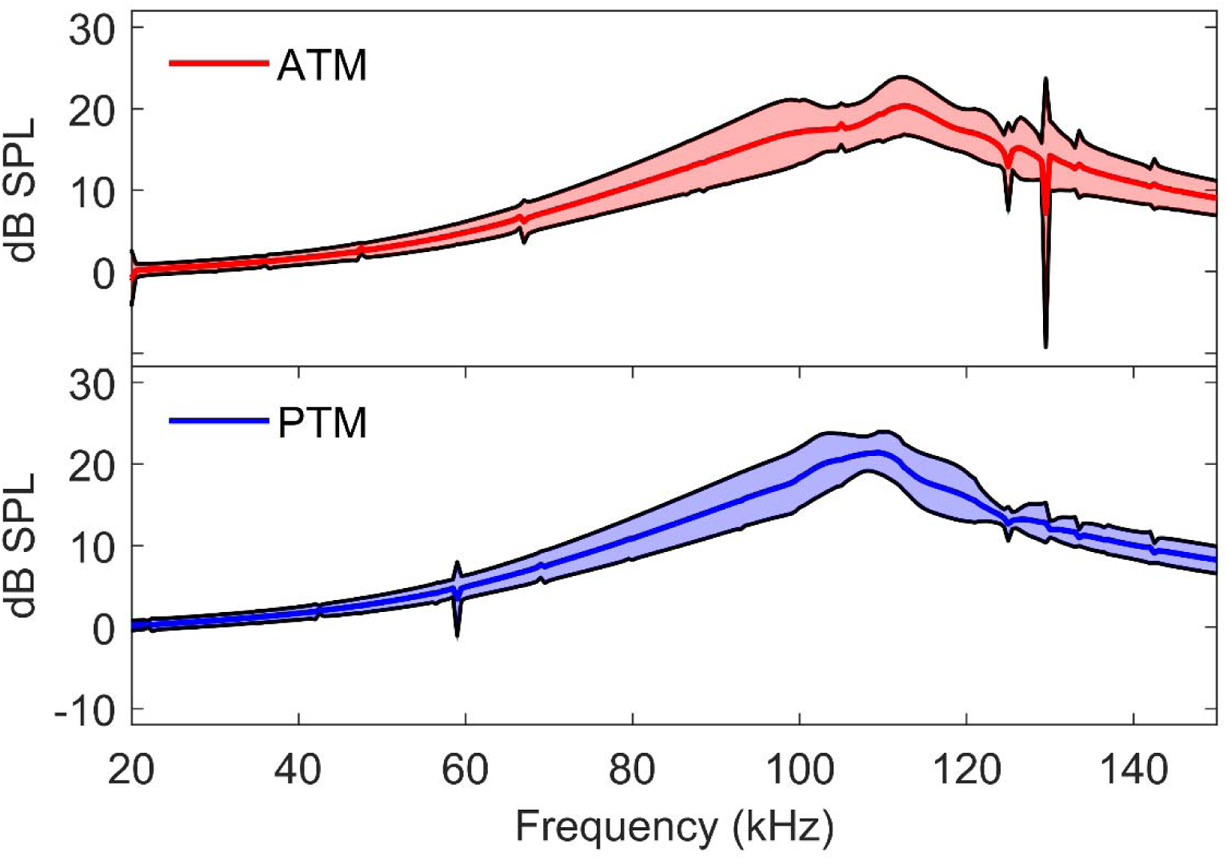
Simulated responses to scaled broadband stimuli incorporating 3D print model ear material properties (dB_peak_ at ∼ 105 kHz for both ATM and PTM). ATM in red bars and PTM in blue bars.

**Figure S3.**
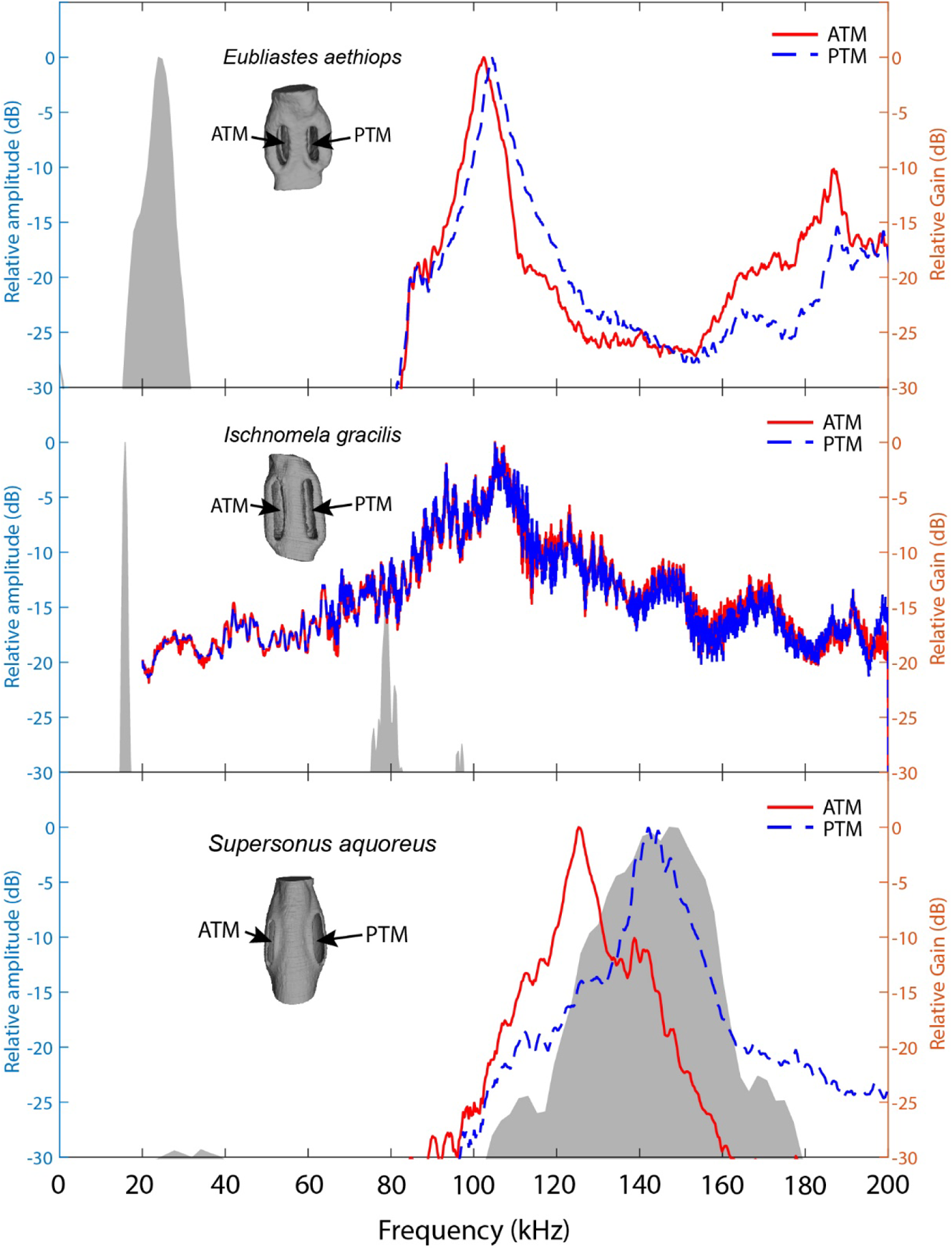
Tympanal cavity relative gain (dB) of sympatric katydid species with auditory pinnae, from the island of Gorgona, Colombia, in response to scaled broadband chirps. *Ischnomela gracilis* (Pseudophyllinae) printed scale 1:12 showing resonance peaks of 104 kHz in both ATM and PTM, *Eubliastes aethiops* (Pseudophyllinae) printed scale of 1:13.1 showing resonance peaks of 102 kHz ATM and 104 kHz PTM, and *Supersonus aquoreus* (Meconematinae) printed scale 1:21.5 showing resonance peaks of 125 kHz ATM and 142 kHz PTM. ATM in red lines and PTM in blue lines. Shaded grey area represents a frequency spectrum of the conspecific calling song of each species (relative amplitude dB).

**Figure S4.**
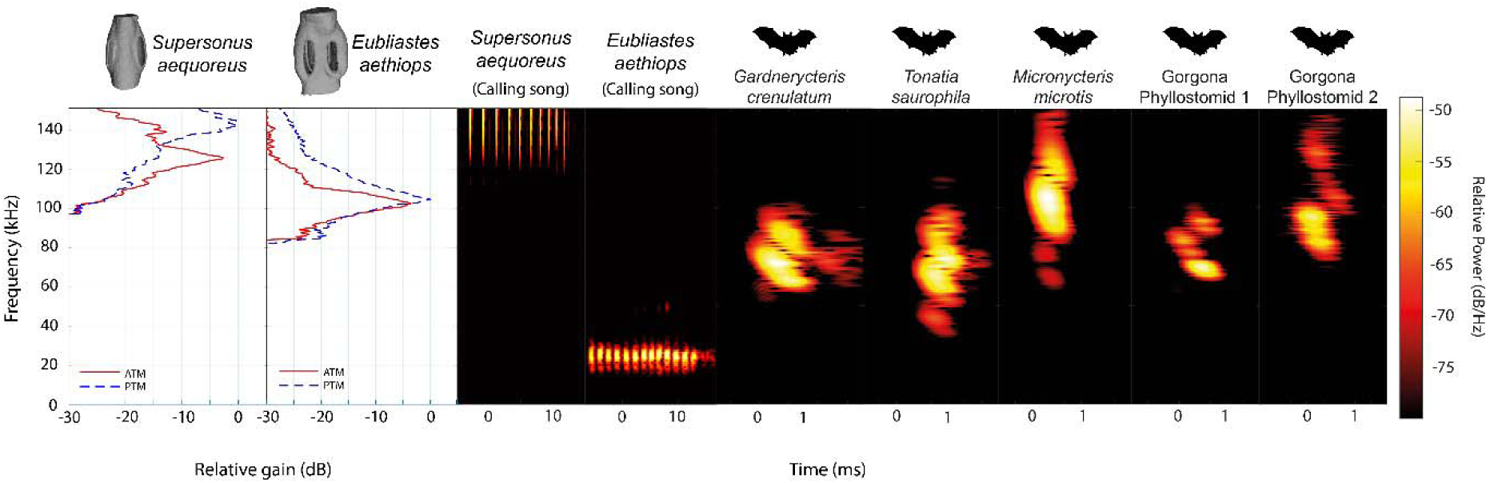
Ecological relevance of pinnae in *Supersonus aequoreus* and *Eubliastes aethiops*. Sound pressure level gains (left) induced by the pinnae are present only at frequencies >80 kHz, covering the range of echolocation frequencies of the three native insectivorous gleaning bat species recorded for Gorgona (10) Gorgona Phyllostomid 1 and Gorgona Phyllostomid 2 represent two examplary bat calls recorded on the island. The conspecific call of *Eubliastes aethiops* is not enhanced by the pinnae (dB_peak_ at 22 kHz); however *Supersonus aequoreus* likely uses pinnae for both conspecific calls and bat detection, if predated by bats (dB_peak_ at 144 kHz). Spectrogram parameters: FFT size 512, Hamming window, 50% overlap, frequency resolution: 512 Hz, temporal resolution: 0.078 ms.

**Figure S5.**
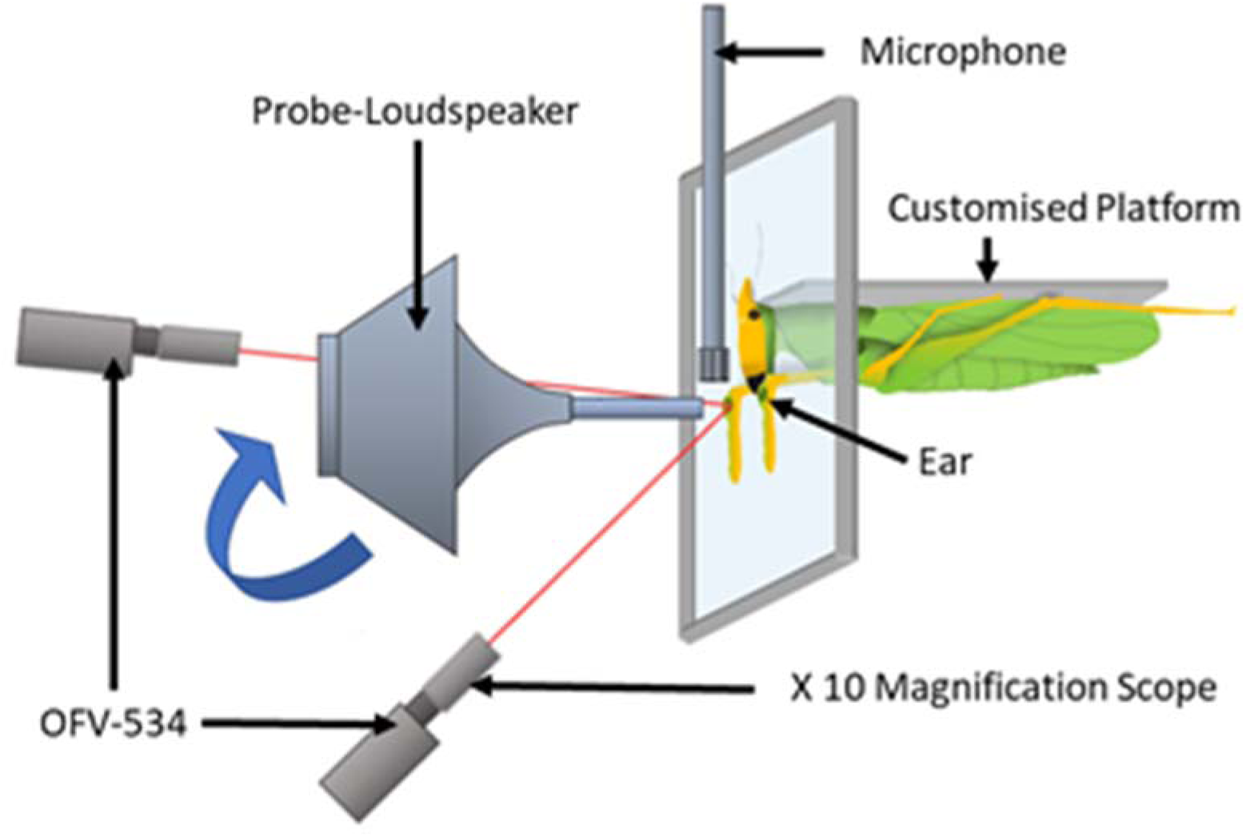
Diagram of experimental arena with a mounted *Copiphora gorgonensis* (not drawn to scale) on specialised (input isolating) platform with a rotating probe-tipped loudspeaker perpendicular to the tympanal septum and single point sensor heads (OFV-534) in perpendicular position.

**Figure S6.**
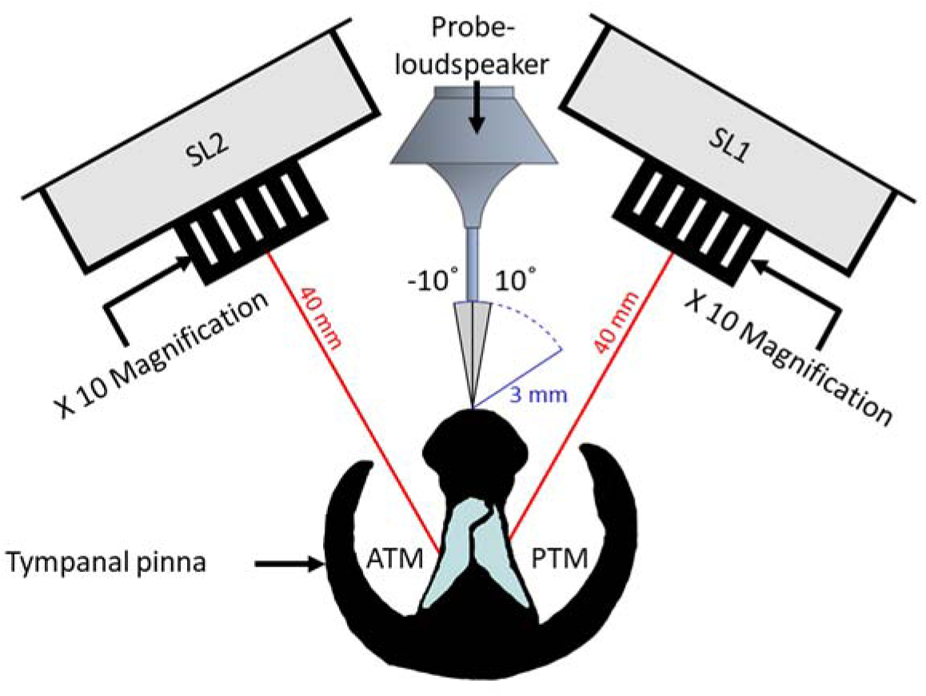
Illustration of experimental arena with a cross section of the copiphorine ear positioned in relation to the single point laser sensor heads with magnifying lenses (SL1 and SL2) and the rotating probe-loudspeaker. The loudspeaker was rotated along a 12 cm semi-circle in 1° steps (each automated step about 0.56 mm) from −10° to 10° perpendicular to the centre of the ear and presented a four-cycle sinusoidal wave at the frequencies 23, 40 and 60 kHz in sequence.

**Figure S7.**
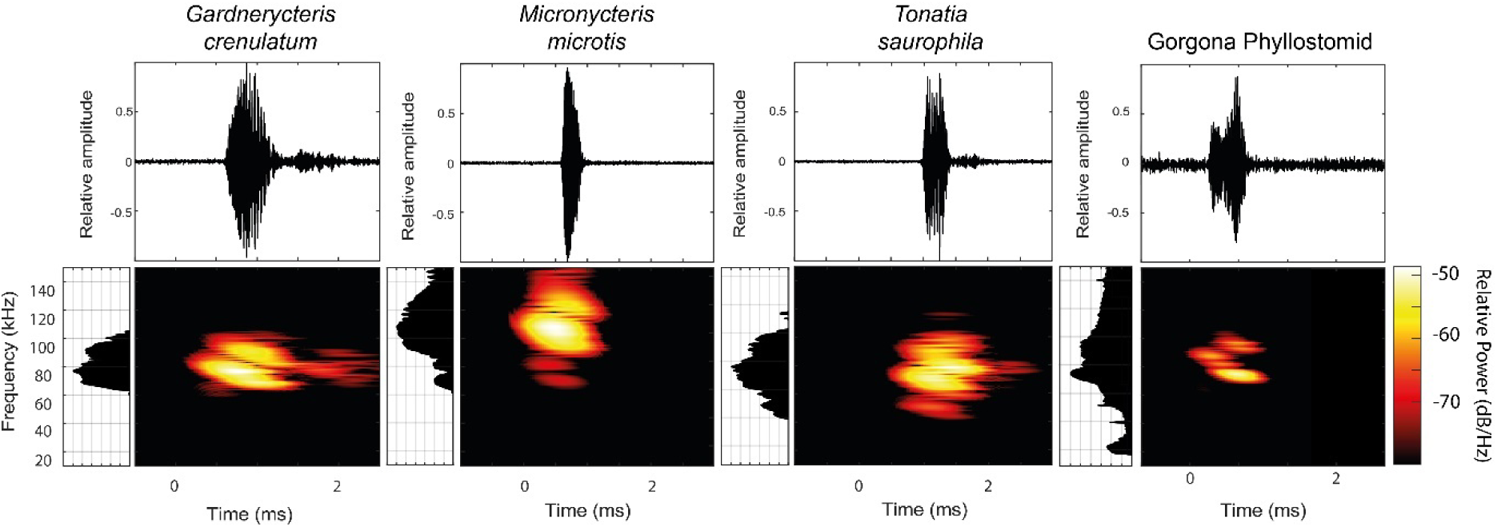
Elaborated echolocation descriptions from four (*Gardnercyteris crenulaum, Micronycteris microtis, Tonatia saurophila*, Gorgona Phyllostomid) insectiverous phyllostomid bats (Chiroptera: Phyllostomidae). For each species, the waveform (top), frequency spectrum and FFT power spectrum (lateral to each spectrogram, 30 dB below peak shown) of a single call is presented. Spectrogram parameters: FFT size 512, Hamming window, 50% overlap, frequency resolution: 512 Hz, temporal resolution: 0.078 ms.

**Figure S8.**
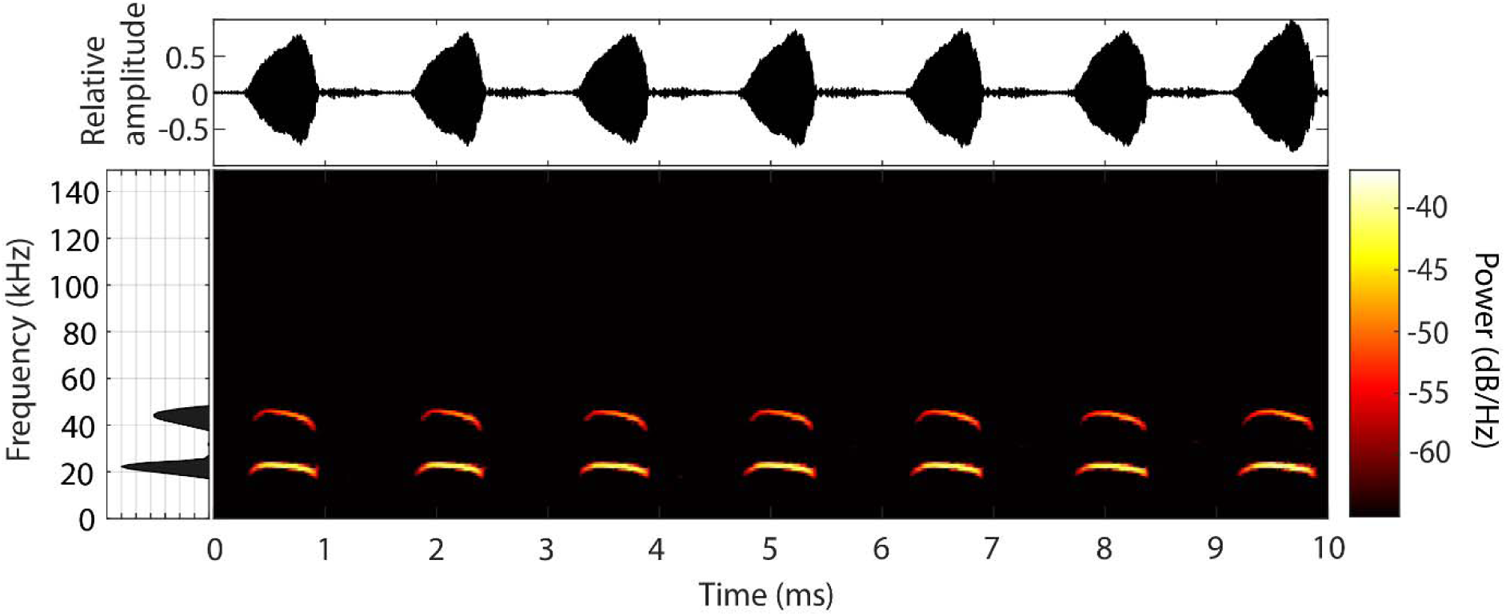
Elaborated conspecific call description of male *Copiphora gorgonensis*. Waveform of multiple syllables within a call, showing the interval between chirps (top). Spectrogram of full signal in the same time range, and FFT power spectrum (lateral, 30 dB below peak shown). Spectrogram parameters: FFT size 512, Hamming window, 50% overlap, frequency resolution: 512 Hz, temporal resolution: 0.078 ms.

**Figure S9.**
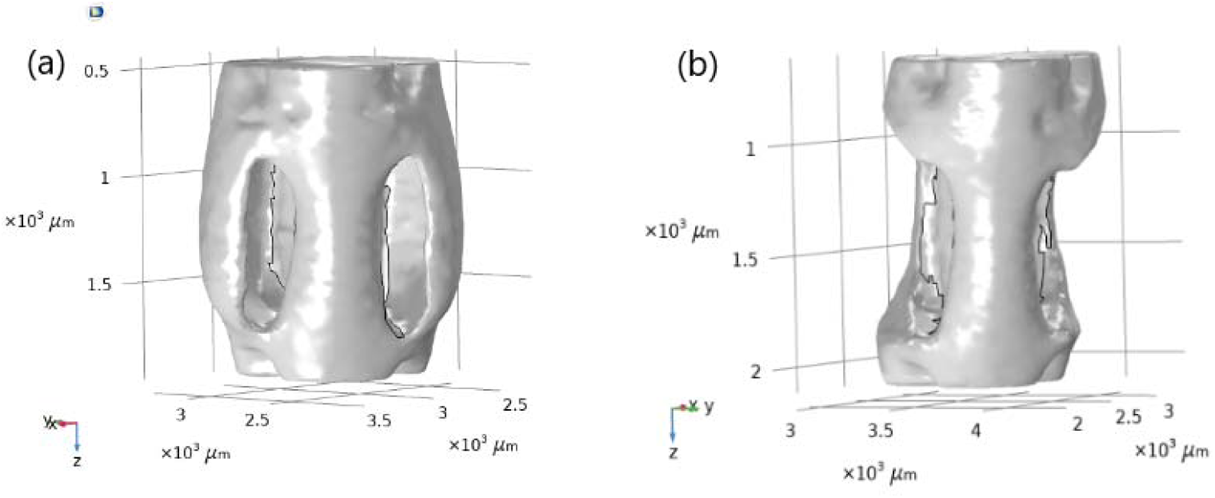
Ear geometries of (a) pinnae intact and (b) pinnae ablated used in the mathematical model for frequency and time domain calculations.

**Figure S10.**
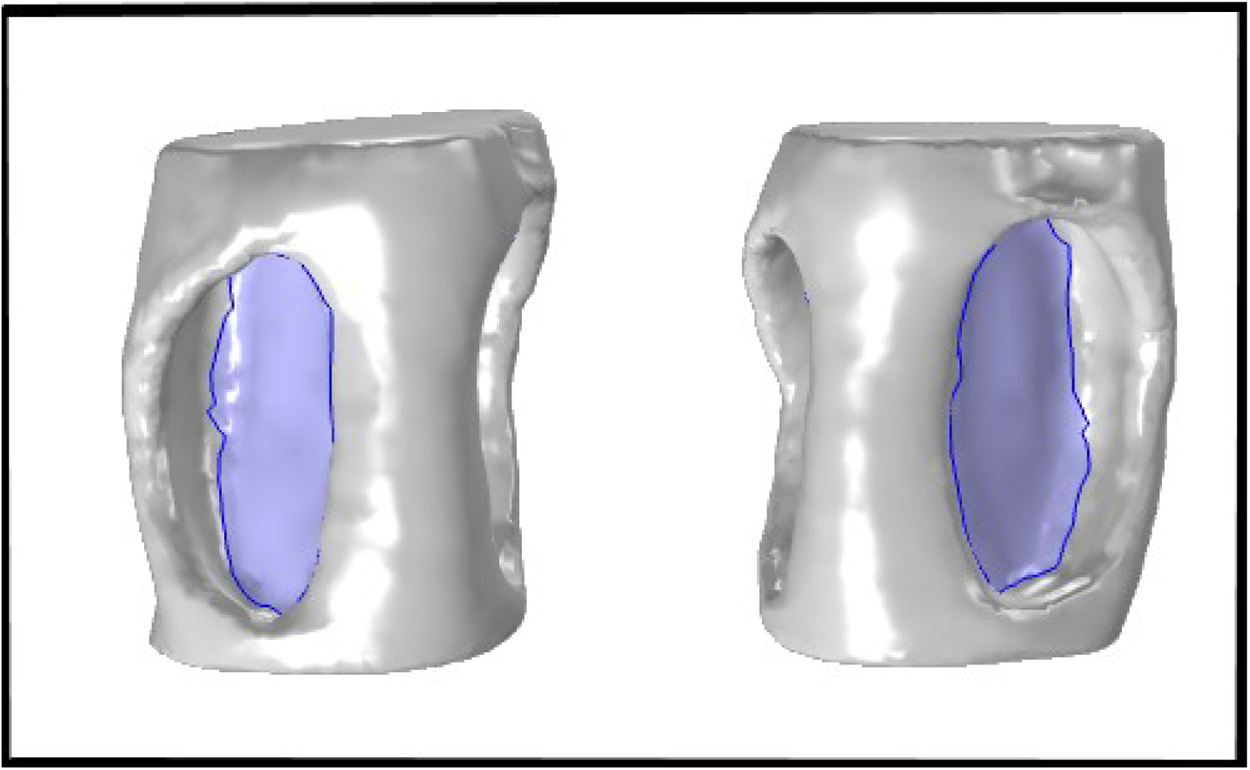
The ear as an isotropic shell system used for the calculation of displacement and stresses resulting from the fluid load. The tympanic membranes were defined as a shell made of a linear, elastic material with a Young’s modulus of 2 GPa, density of 1300 kg/*m*^3^, Poisson’s ratio of 0.3, and thickness 5 μm (11).

**Figure S11.**
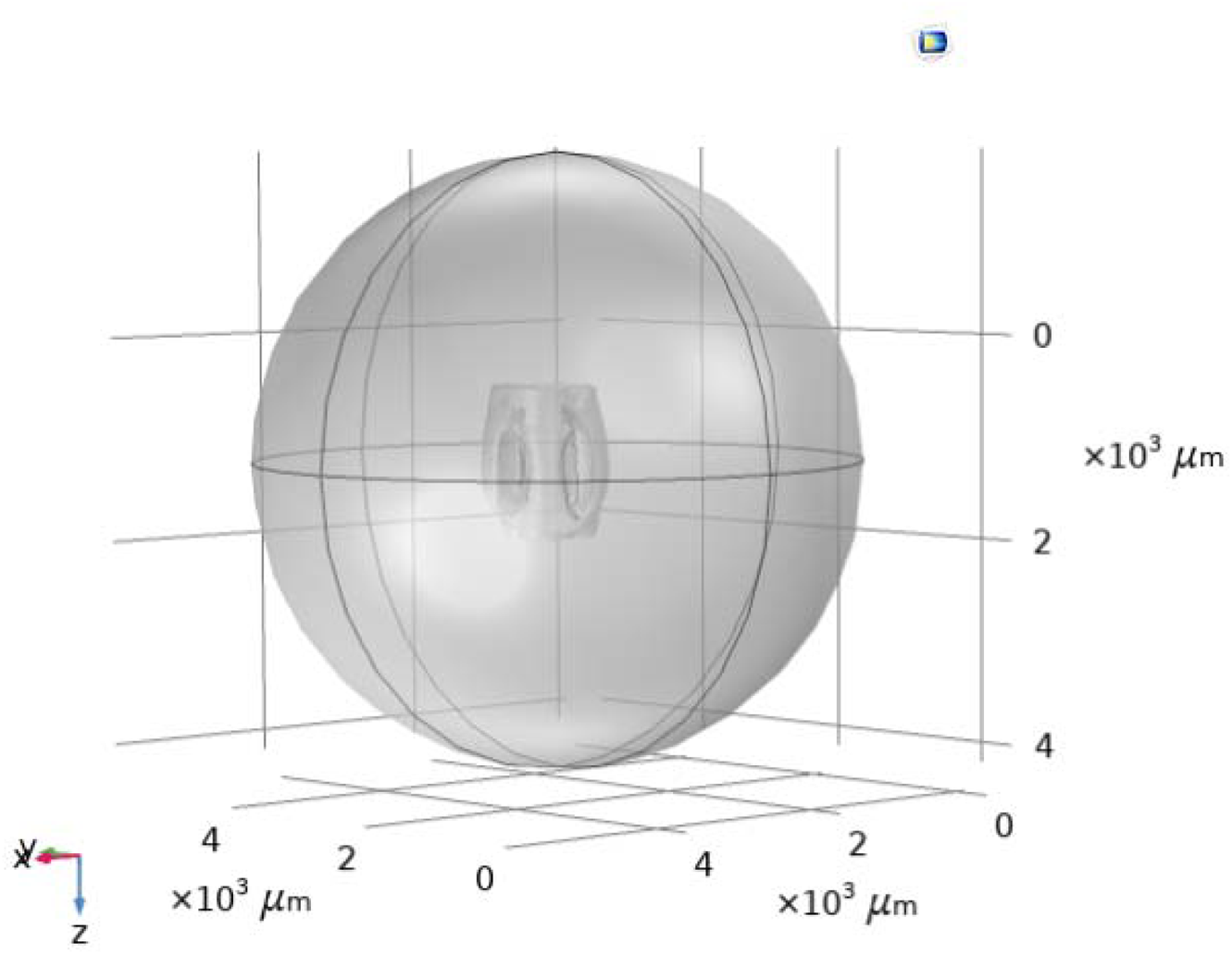
The air acoustic domain of a truncated sphere with a 3 mm radius centered around the reconstructed micro-CT geometry of the ear.

**Figure S12.**
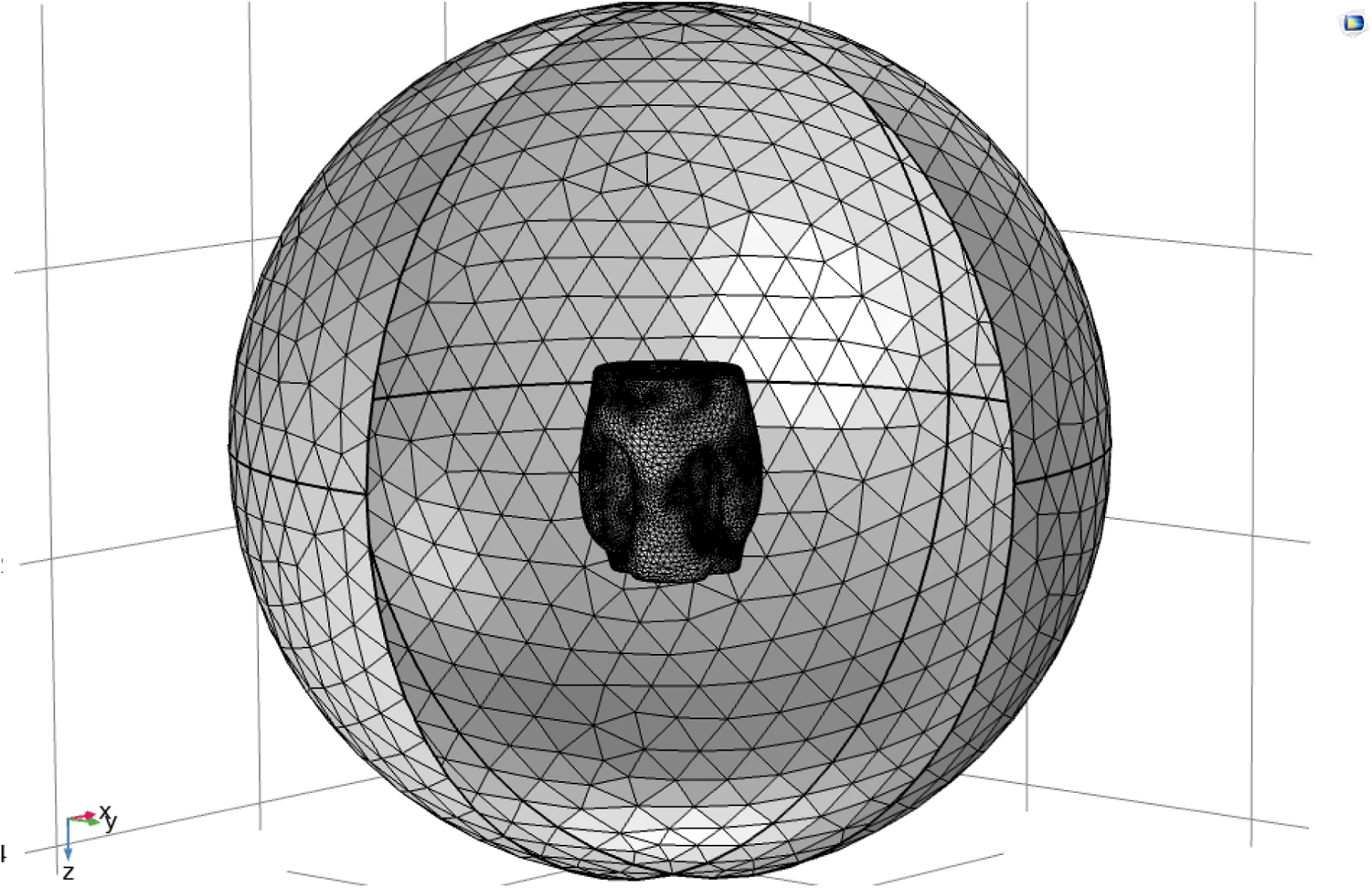
Finite element mesh sphere composed of tetrahedral elements of maximum diameter 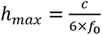, where *c* = 343 m/s and *f*_0_ = 150 kHz enclosing 3 mm diameter ear.

**Figure S13.**
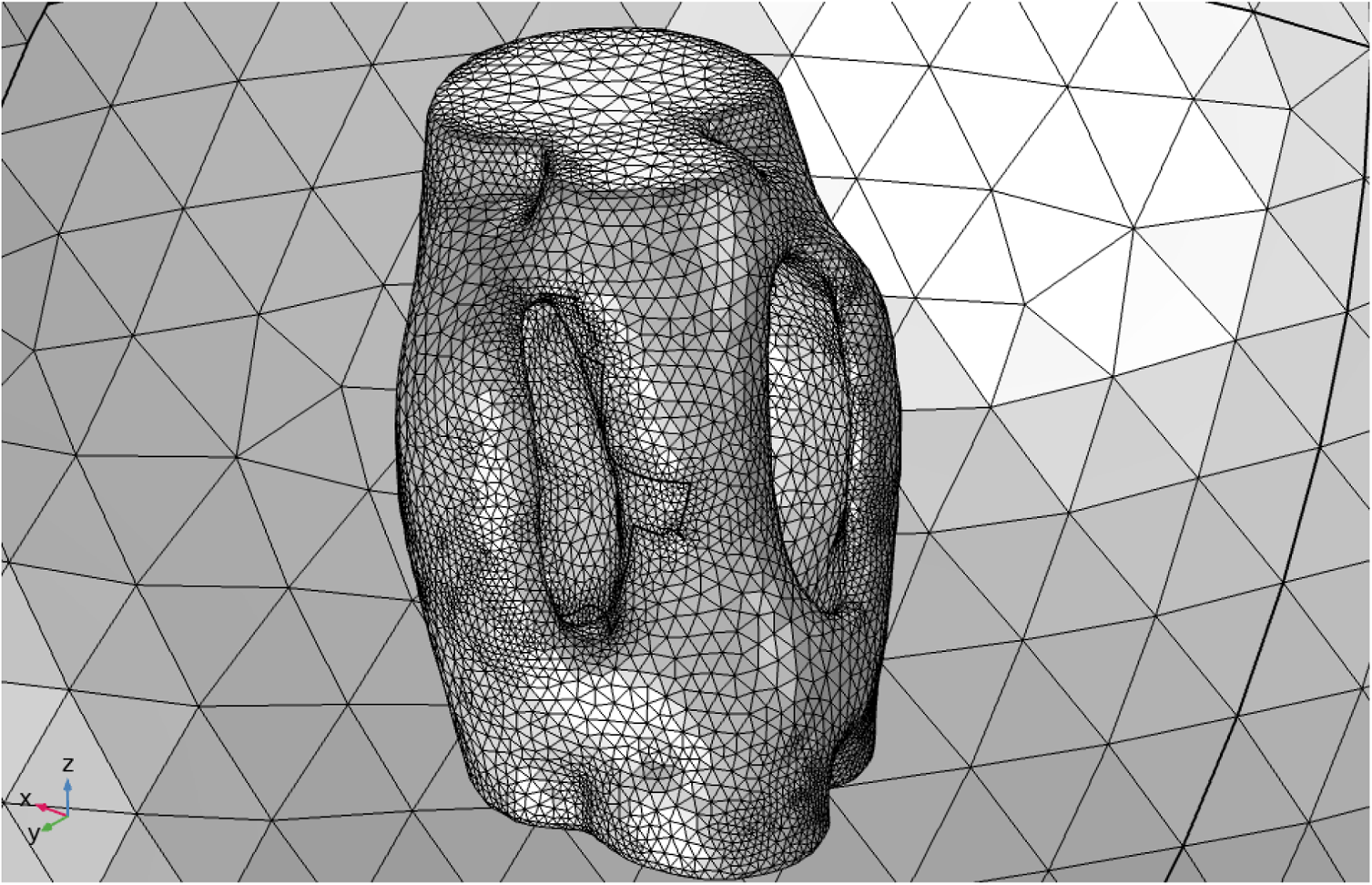
An enlarged diagram (from fig. S12) of the insect ear with finite element mesh.

**Table S1.**
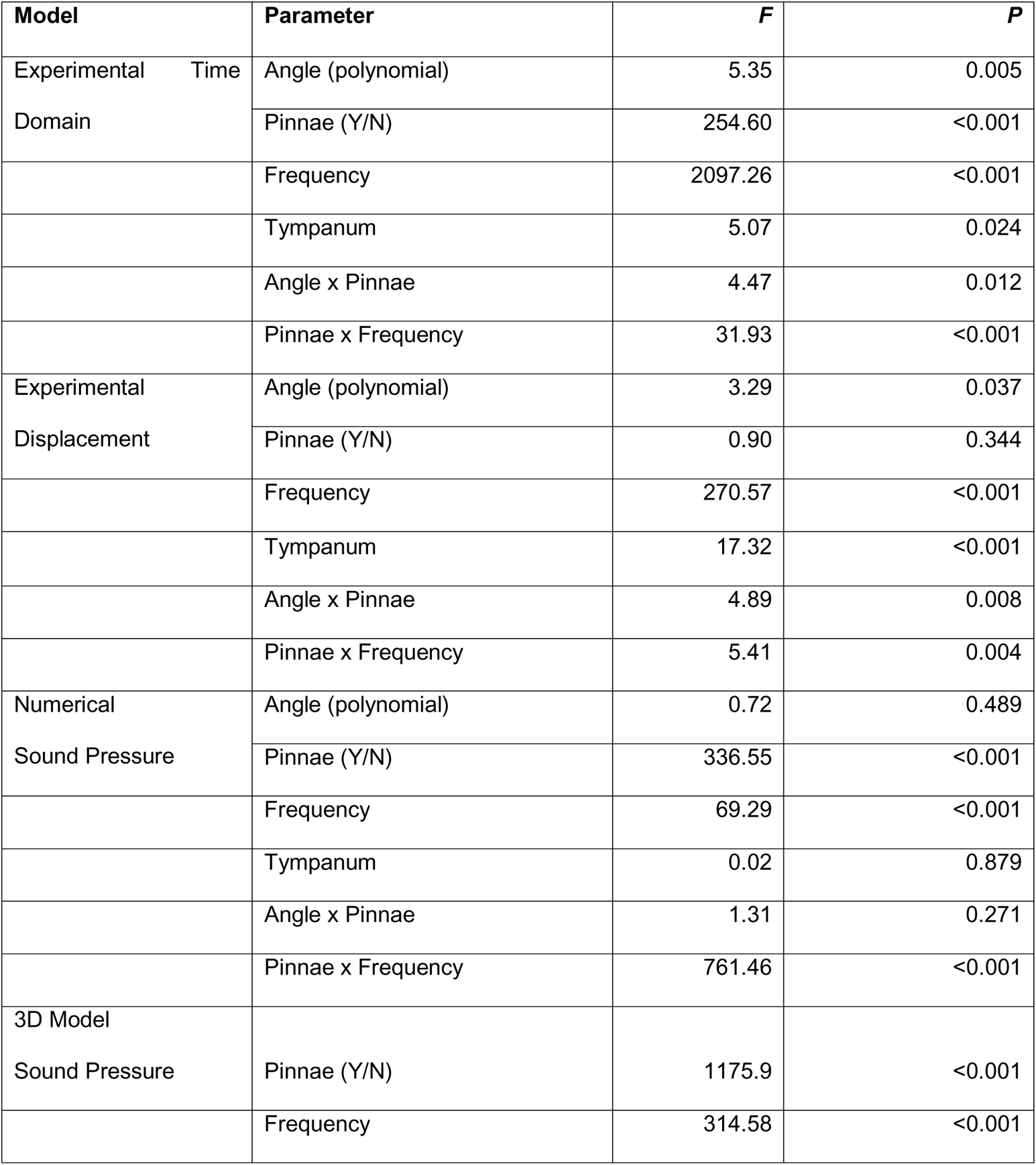
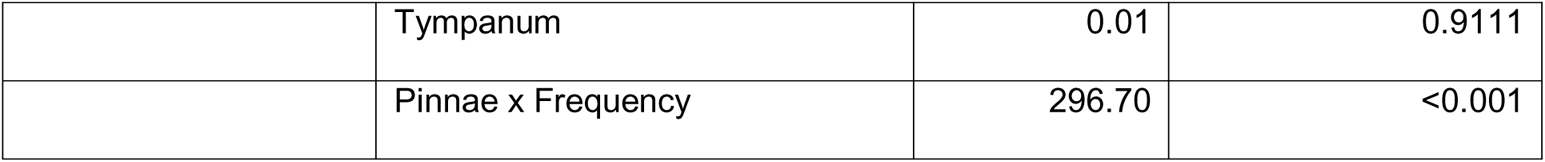
Linear mixed models (LMM) of experimental and numerical simulation data. Parameters showing effects of angle, pinnae, frequency, tympanum, angle x pinnae and pinnae x frequency for time domain data (experimental time and displacement) and sound pressure (numerical simulations and 3D print models). Experimental models n = 13 ears. Numerical model n = 17 ears. 3D model n = 4 ears.

**Table S2.** Measured parameters of the ear of C. gorgonensis (n = 8 ears; 3 females, 2 males). Given are mean values (± SD). Abbreviations: ATM = anterior tympanal membrane; PTM = posterior tympanal membrane.

| ATM<br>cavity<br>volume<br>(mm<sup>3</sup>) | PTM<br>cavity<br>volume<br>(mm<sup>3</sup>) | Distance<br>between<br>slits (mm) | Cross-sectional<br>width of<br>foreleg<br>below ear<br>(mm) | Cross-sectional<br>width of<br>ear (mm) | ATM<br>slit<br>area<br>(mm<sup>2</sup>) | PTM<br>slit<br>area<br>(mm<sup>2</sup>) | Protrusion<br>of anterior<br>pinna (mm) | Protrusion<br>of posterior<br>pinna (mm) |
| --- | --- | --- | --- | --- | --- | --- | --- | --- |
| 0.14<br>($\pm 0.01$) | 0.15<br>($\pm 0.01$) | 0.42<br>($\pm 0.03$) | 0.84<br>($\pm 0.02$) | 1.14<br>($\pm 0.35$) | 0.16<br>($\pm 0.01$) | 0.16<br>($\pm 0.01$) | 0.39<br>($\pm 0.02$) | 0.45<br>($\pm 0.03$) |

**Table S3.** Raw data for the behavioural audiogram of ultrasound response in nine female *C*. *gorgonensis*. NaN denotes that no response was shown to a particular stimulus. Mean and standard deviation calculated ignoring missing data (NaN) for each frequency in the lower rows of the table. All values in dB SPL.

| ID | Frequency (kHz) |  |  |  |  |  |  |  |  |  |  |  |  |  |  |  |  |  |  |  |  |
| --- | --- | --- | --- | --- | --- | --- | --- | --- | --- | --- | --- | --- | --- | --- | --- | --- | --- | --- | --- | --- | --- |
|  | 20 | 25 | 30 | 35 | 40 | 45 | 50 | 55 | 60 | 65 | 70 | 75 | 80 | 85 | 90 | 95 | 100 | 105 | 110 | 115 | 120 |
| F1 | 90 | 55 | NaN | 50 | NaN | 55 | 50 | 50 | NaN | NaN | NaN | NaN | 55 | NaN | 55 | NaN | 50 | 50 | 55 | 50 | 60 |
| F2 | 85 | 60 | 55 | 55 | 55 | 50 | 55 | 55 | 55 | 55 | 55 | NaN | 60 | 55 | 60 | 55 | 55 | 55 | 55 | 55 | 55 |
| F3 | 80 | 60 | 60 | 60 | 60 | 60 | 60 | 60 | 60 | 60 | 60 | 60 | 60 | 60 | NaN | 60 | NaN | 60 | 60 | 55 | 60 |
| F4 | 75 | 65 | 65 | 55 | 65 | 65 | 65 | 65 | 65 | 65 | 65 | 65 | 65 | 65 | 65 | 65 | 70 | NaN | 60 | 65 | NaN |
| F5 | 60 | 70 | 50 | 50 | 70 | 70 | 50 | 70 | 60 | 70 | 70 | 55 | 50 | 70 | NaN | NaN | NaN | 55 | 75 | 50 | 70 |
| F6 | 75 | 75 | 55 | 50 | 50 | 60 | NaN | NaN | 60 | 55 | 80 | 75 | NaN | 50 | 60 | NaN | 60 | NaN | 60 | 70 | 50 |
| F7 | 80 | 70 | 60 | 50 | 55 | 55 | 50 | 60 | 55 | 55 | 50 | 50 | 50 | 50 | 50 | 65 | 65 | 55 | NaN | 60 | 55 |
| F8 | 85 | 55 | 55 | 55 | 50 | 50 | NaN | NaN | 55 | 60 | NaN | 50 | NaN | NaN | 55 | 50 | 70 | 65 | 50 | NaN | NaN |
| F9 | 90 | 60 | 60 | 50 | 55 | 60 | 55 | 50 | 60 | NaN | 55 | NaN | 60 | 60 | 55 | 55 | 50 | 50 | NaN | 60 | 55 |
| Mean | 80 | 63 | 57.5 | 53 | 57.5 | 58 | 55 | 59 | 58.8 | 60 | 62.1 | 59 | 57 | 58.6 | 57.1 | 58.3 | 60 | 56 | 59 | 58 | 58 |
| STD | 9 | 7 | 4.63 | 4 | 7.07 | 6.6 | 5.77 | 7.5 | 3.54 | 5.77 | 10.4 | 9.7 | 5.7 | 7.48 | 4.88 | 6.06 | 8.7 | 5.3 | 7.9 | 7 | 6.4 |

