## Supplemental Index for "Ear pinnae in a neotropical katydid (Orthoptera: Tettigoniidae) function as ultrasound guides for bat detection"

**This PDF file includes:**

**Supplemental Materials and Methods Text**

**Videos V1 to V3**

**Supplementary Materials and Methods Text**

***Live experiments***

The original generation of the species were imported to the UK under the research permit granted by the Colombian Authority (DTS0-G-090 14/08/2014) in 2015. The specimens were ninth generation, captive bred colonies and maintained at 25°C, 70% RH, light: day 11 h: 23 h. They were fed *ad libitum* diet of bee pollen (Sevenhills, Wakefield, UK), fresh apple, dog food (Pedigree Schmackos, UK) and had access to water.

***Insect call recordings***

Sound recordings were performed in a sound-attenuated booth at the Sensory Biology Lab, University of Lincoln, UK, at a temperature of 25ºC and relative humidity of 40%. The specimens were placed on a metallic screen cage at 10 cm from a 1/8” microphone (B&K Type 4138 omnidirectional microphone, Brüel & Kjær, Nærum, Denmark), connected to a 1/4” preamplifier (B&K 2670, Brüel & Kjær, Nærum, Denmark) and set to a conditioning amplifier (Nexus 2690-OS1, Brüel & Kjær, Nærum, Denmark). The microphone was calibrated at 94 dB SPL (re 20 µPa), using a B&K sound level calibrator (Type 4231, Brüel and Kjaer, Nærum, Denmark). Data was obtained via an acquisition board (PCI-6110, National Instruments, Austin, TX, USA) and stored on a computer hard disk at a sampling rate of 512 kHz using the Polytec acquisition software (PSV 9.0.2, Polytec GmbH, Waldbronn, Germany). Sound was analyzed using Matlab (R2015a, The MathWorks, Inc., Natick, MA,USA) (*SI Appendix*, Figure S4).

To 3D print the ears of *Copiphora gorgonensis*, *Ischnomela gracilis*, *Supersonus aquoreus* and *Eubliastes aethiops*, micro-CT stereolithography files (STL) were imported into the software CHITUBOX 64 (Chitubox, Guangdong, China). The models were scaled to be approximately 12x larger than the actual ears. Support structures and a base printing platform were then added to support the model, with a 0.2 mm attachment thickness to the model. Supported models were delivered via USB to a Mars Elegoo Pro 2 3D Printer (Elegoo Inc., Shenzhen, China). Models were printed using grey ABS-like photopolymer resin (exposure parameters: 20 s first layer, 5 s normal layers) with a solidification wavelength of 405 nm. When printing was complete (about 1 h 30 min), models were washed in 100% isopropyl alcohol, rinsed in cold water, then exposed to UV light in an Elegoo Mercury Plus curing station (Elegoo Inc., Shenzhen, China) for 8 min. To prepare the models for entry of the probe microphone into the tympanal cavities, 2 mm diameter holes were drilled into the centre of the base of each cavity (Fig. 3).

***Mathematical models and numerical simulations methods***

For the numerical simulation of the problem, we solved a system of equations representing the sound pressure (SPL) inside and around the *C. gorgonensis* ear, resulting from the interaction of the ear with an incident plane acoustic wave in an air domain. The air acoustic domain is truncated as a sphere with a 3 mm radius that is centered around the ear (*SI Appendix*, Fig. S10).

Two different sets of mathematical models were considered in the described geometry, within the frequency and the time domains. For the frequency domain calculations, the solution to the Helmholtz equation

$$\frac{1}{\rho}\Delta p_{f}+k^{2}p_{f}=0 (1)$$

was considered for the acoustic system, where the parameters $\rho$ = is the density of air, $k=\omega/c$ is the wavenumber, $\omega$ is the angular frequency and $c$ = 343 m/s is the speed of sound in air. The variable $p_{f}\left( x \right)$ is the total pressure in the frequency domain, which is dependent on the 3D spatial variables $x=\left( x,y,z \right)$, and $\Delta=\frac{\partial^{2}}{{\partial x}^{2}}+\frac{\partial^{2}}{{\partial y}^{2}}+\frac{\partial^{2}}{{\partial z}^{2}}$ is the Laplace operator.

At the outer perimeter of the sphere, to allow for a radiated or scattered spherical wave to travel out of the modelling domain without reflections, a spherical radiation boundary condition was applied in the following form:

$$n.\nabla p_{f}+\left( ik+\frac{1}{r} \right)p_{f}-\frac{r\Delta_{||}p_{f}}{2\left( ikr+1 \right)}=n. \nabla p_{fi}+\left( ik+\frac{1}{r} \right)p_{fi}-\frac{r\Delta_{||}p_{fi}}{2\left( ikr+1 \right)}, (2)$$

where $n$ is the normal vector, *r* is the distance from the source location, the operator $\Delta_{||}$ denotes the Laplace operator in the tangent plane at a particular point and $i=\sqrt{-1}$. This boundary condition was based on an expansion in spherical coordinates given in Bayliss *et al*. (Bayliss et al., 1982) and implemented to the second order. The right-hand side of equation (2) allows for an incoming plane wave defined as

$$p_{fi}=e^{-ik\left( \frac{x.e_{k}}{||e_{k}||} \right)}$$

with magnitude 1 Pa and frequency ranging from 2 to 150 kHz. The wave travels from the direction $e_{k}$, which was taken as normal to the front of the ear (“point zero”).

The ear itself was considered as an isotropic shell system which allowed for the calculation of displacement and stresses resulting from the fluid load. The tympanic membranes were defined as a shell made of a homogeneous, linear elastic material with a Young’s modulus of 2 GPa, density of 1300 kg/$m^{3},$ Poisson’s ratio of 0.3, and thickness 5 μm (Montealegre-Z and Robert, 2015) (*SI Appendix*, Fig. S10). The rest of the ear was assumed to have a thickness of 175 μm and the same material properties as the tympana.

Finally, the continuity between the acoustic and shell systems was retained by accounting for the interaction between the two systems. After calculating the frequency response of the ear to the fluid load in the form of harmonic displacements and stresses, the model used the displacement magnitude of the solid surface in the acoustic domain inner boundary to ensure continuity. This is represented by the equations

$$n.\frac{1}{\rho}\nabla p_{f}=\omega^{2}U_{sf},$$

$$F_{Af}=p_{f}n,$$

At the intersection of the ear with the sphere, where $U_{sf}$ is the ear (shell) displacement and $F_{Af}$ is the load (force per unit area) experienced by the shell structure.

An analogous model was also considered in the time domain, for which instead of equation (1), the wave equation

$$c^{2}\Delta p_{t}=\frac{\partial^{2}p_{t}}{\partial t^{2}}$$

was solved for in the acoustic domain, where $p_{t}(x,t)$ is the total pressure in the time domain, which is dependent on both the space variables $x$ and the time variable $t$. The boundary condition (2) was also replaced by the time dependent spherical wave condition

$$n.\nabla p_{t}+\left( \frac{1}{c}\frac{\partial p_{t}}{\partial t}+\frac{1}{r}p_{t} \right)=n. \nabla p_{ti}+\left( \frac{1}{c}\frac{\partial p_{ti}}{\partial t}+\frac{1}{r}p_{ti} \right),$$

where the incident wave $p_{ti}=\sin\left( 2\pi f_{0}\left( t-\frac{x.e_{k}}{c||e_{k}||} \right) \right)$, at frequencies $f_{0}=23, 40 \mathrm{and} 60 \mathrm{kHz}$. For the time domain models, five different incident wave direction were considered, which were -10°, -5°, 0°, 5° and 10° on a fixed plane perpendicular to the ear, where 0° is analogous to the “point zero” defined during the section *vibration measurements* in the main text.

Finally, the continuity of the acoustic and shell systems was ensured with the equations

$$n.\frac{1}{\rho}\nabla p_{t}=\frac{\partial U_{st}^{2}}{\partial t^{2}},$$

$$F_{At}=p_{t}n,$$

at the intersection of the ear with the sphere, where $U_{st}$ is the time dependent displacement of the ear and $F_{At}$ is the time dependent load experienced by the shell structure.

The numerical solution to the problem was obtained using the finite element method for the spatial variables in both the time and frequency domain simulations. For forming the finite-element mesh, the maximum diameter used for the tetrahedral elements in the sphere was $h_{max}=\frac{c}{6\times f_{0}}$, where $c=343 m/s$ and $f_{0}=150 \mathrm{kHz}$ (*SI Appendix*, Fig. S12 and S13). Hence, even at the largest frequency considered, there were six tetrahedral elements per wavelength. Quadratic Lagrange elements were applied for the finite element solution. For the time domain solution, the time variable was solved for using the Generalized alpha method, with a constant time step of $\Delta t=\frac{1}{60\times150}s,$ so that the Courant-Friedrichs-Lewy (CFL) condition (Courant et al., 1967), defined as $CFL= \frac{c\times h_{max}}{\Delta t}$ was 0.1, which gives a reliable approximation of the solution.

***Behavioural audiograms***

For behavioural audiograms nine females were tethered from the pronotum to control for a constant position sound pressure, while the specimen walked on a foam rotating cylinder. The cylinder was 15 cm diameter by 15 cm depth, and was customised by the Foam Superstore (<https://foamsuperstore.co.uk/foam-circles-cylinders-cut-to-size.html>). The cylinder freely rotated on a rod crossing along its longitudinal axis, with each end resting on the centre of a Hard Disk Drive Spindle Wheel (custom designed using parts of old computer hard drives). These wheels produce smooth rotation of the rod and cylinder that do not disturb the insect. Specimens were glued from the pronotum to a 25 cm wooden rod (4 mm diameter) using bees wax (Fisher Scientific UK, Limited, Leicestershire; product code W/0200/50) and Colophony resin (Sigma-Aldrich Co. St. Louis, MO, USA; Product No. 60895-250G) in a 1:1 mix . The wooden rod was held by a micromanipulator which allows positioning of the insect on the rotating foam cylinder. Each specimen was left to adapt to the new situation for 15 minutes, before the experiment started. This experimental setup was mounted on a Pneumatic Vibration Isolation Table (B120150B - Nexus Breadboard, 1200 mm x 1500 mm x 110 mm, Thorlabs Inc., USA) supported by an anti-vibration frame (PFA52507 - 800 mm Active Isolation Frame 900 mm x 1200 mm, Thorlabs Inc., USA). All experiments were conducted inside an acoustic booth (AC Acoustics, Series 120a, internal dimensions of 2.8 m x 2.7 m x 2.7 m). A disadvantage of the treadmill used here was that the insect is forced to walk in the forward direction, different to other more sophisticated air-cushioned spherical treadmill systems that allow movement in any direction (Hedwig and Poulet, 2004; Mason et al., 2001). However, since we were not interested in directional responses, but only on startle behaviour, this simple treadmill was useful.

Acoustic stimuli were generated in a function generator (Agilent 33120A, 15MHz Function/Arbitrary waveform generator, Agilent Technologies UK Ltd., Edinburgh, UK), and shaped into 10-ms pulses (2-ms linear rise/fall) at 6 volts peak-to-peak. Function generator output was connected into a portable single channel ultrasonic power amplifier suited for the ultrasonic speakers, Model B without 200V bias voltage generator (Avisoft Bioacoustics, Glienicke/Nordbahn, Germany). Sound stimulus was delivered using a SS-TW100ED Super-Tweeter loudspeaker (Sony, Tokyo, Japan), which has a frequency response in the range 20 to 125 kHz. The input from the Avisoft amplifier was high-pass filtered at 20 kHz using the built-in filter of the Sony Tweeter. The speaker was positioned 15 cm antero-lateral from the specimen. The amplitude of the stimulus was monitored using a 1/8” precision pressure microphone (Bruel & Kjaer, 4138; Nærum, Denmark) and a preamplifier (Bruel & Kjaer, 2633; Nærum; Denmark). The microphone was calibrated using a sound level calibrator (Bruel & Kjaer, 4231; Nærum, Denmark), and positioned 5 cm above of the tethered insect. The acoustic stimuli were constantly monitored in real time using the analyser window of the Polytec laser software (Polytec; Waldbronn, Germany).

At each frequency (20 : 5 : 120 kHz), pure tones of 10 ms duration were played at increasing amplitude (40 : 5 : 90 dB SPL) to measure behavioural thresholds of the nine female *C. gorgonensis*. Starting at the lowest amplitude for a given frequency, each stimulus lasted 1 s, and consisted of ten 10-ms pulses presented at a rate of 100 Hz. Three types of behaviours were observed: (1) interruption of walking; (2) alert (the katydid tried to jump or adopted a defensive position); (3) no response. If any of reactions (1 and 2) occurred, the stimulus was decreased by 5 dB and the animal was re-tested once walking resumed. Threshold was defined as the lowest amplitude that reliably elicited a behaviour and for each given frequency. We anticipated that above this sound pressure, the insect continued hearing the stimulus. If no response occurred, the stimulus was repeated (after a few seconds of silence) to verify the lack of response. If still no response, the stimulus amplitude was increased by 10 dB steps and the katydid was re-tested.

For purposes of analysis, for each specimen the threshold at each frequency was annotated in a matrix for further calculation of mean vector and standard deviations. Not all specimens showed consistent response at all frequencies and treatments, and if no response was shown to a particular stimulus, but the specimen was shown response to other stimuli, the missing response was entered as NaN (missing value identifier for Matlab matrix computation; see Supplementary Table S3).

***Neural audiograms***

Suction electrode recordings were obtained from the auditory nerves of five adult *C. gorgonensis* following previously described methods (Isaacson and Hedwig, 2017). Briefly, animals were restrained dorsal side up in plasticine with their acoustic spiracles and tympana exposed to the air. One auditory nerve was sampled per animal, which was accessed by removing a small window of cuticle from a front femur and dissecting away any obstructing material. A pre-prepared polycarbonate electrode (1 mm outer diameter; 0.5 mm internal diameter; pulled by hand over a soldering iron and cut to a terminal internal aperture of ~40 µm) was filled with HEPES-buffered saline that had been made viscous with 4% Tylose H200 NP2 (ShinEtsu, Wiesbaden, Germany) to prevent leakage from the tip. The electrode was fitted into a custom-made holder, with a platinum wire inserted into the saline. The electrode tip was then placed onto the auditory nerve using a micromanipulator, and sealed using gentle suction. A platinum reference electrode was inserted into a small incision in the distal tibia.

Whole-nerve activity in response to sound was recorded using a differential amplifier (A-M Systems Inc., Carlsborg, WA, USA, model 1700), and sampled at 15 kHz using an analogue-to-digital converter and recording software (CED Micro 1401 and Spike2 version 7, Cambridge Electronic Design, Cambridge, UK). Acoustic stimuli were created by a function generator (SDG1020, Siglent technologies, Augsburg, Germany), consisting of a fully amplitude modulated pulse with a cycle frequency of 1 Hz (0.5 s ON, 0.5 s OFF). This signal was carried via a power amplifier (SA1, Tucker-Davis Technologies System, Alachua, FL, USA) to an ultrasonic power amplifier (Avisoft Bioacoustics, Glienicke/Nordbahn, Germany). From here, the signal passed through a super tweeter (SONY ref) capable of producing acoustic signals from 20 – 125 kHz positioned 15 cm from the animal, with a clear path to both the tympana and acoustic spiracle of the recorded ear. To calibrate the SPL of the signal, a 1/8” condenser microphone (Type 4138, Bruel & Kjaer Nærum, Denmark) with built-in pre-amplifier (Type 2670, Bruel & Kjaer Nærum, Denmark) was connected to the same data acquisition system as the neural recording via a power amplifier (Type 12AA, G.R.A.S., Holte, Denmark). From here, the signal amplitude was calibrated at 94 dB SPL (1 Pa) using a portable sound pressure calibrator (Type 4231, Bruel & Kjaer Nærum, Denmark). The microphone was then placed above the tympanal organ, and the SPL of the stimulus modified until the output SPL was equal to the calibrated 94 dB SPL. To modify the SPL following calibration, the gain output of the aforementioned SA1 power amplifier was reduced in –6 dB steps. Eleven different sound stimuli consisting of pure tones ranging from 23–120 kHz were randomly presented. Each stimulus was presented nine times per frequency at increasing sound pressures from 46–94 dB in 6 dB increments (giving a total of 10 repeats × 9 sound pressures × 11 frequencies = 990 responses per animal).

Individual action potentials from auditory afferents were too small to be individually identified and characterised amidst all the other neuronal activity in the nerve, so the neuronal recordings were root-mean-square (RMS) transformed in Spike2 with a time constant of 0.66 ms to convert the waveform into a displacement from zero. This allowed the neuronal response to sound to be characterised as an area, with units of µVs. An averaged response to each train of ten pulses and succeeding silent periods per sound intensity and frequency was produced in Spike2. The response area to 475 ms of sound stimulus (excluding the transient ‘on’ response immediately after the onset of a sound pulse) and an equivalent 475 ms in the succeeding silent period was measured in each averaged response. The mean area of response when the sound was on was compared to the corresponding mean period of silence for each combination of frequency and sound intensity using paired *t*-tests.

***Statistical analyses***

Using the same initial LMM model, we tested how sound pressure estimated from numerical models was related to angle (-10° to 10°, polynomial continuous variable), presence of pinnae (y/n), frequency (23, 40 and 60 kHz, categorical variable), tympanum (ATM or PTM) as fixed factors. Again, we include the interactions between angle and pinnae and between pinnae and frequency. We finally tested sound pressure based on 3D models with the presence of pinnae (y/n), frequency (23, 40 and 60 kHz, categorical variable), tympanum (ATM or PTM) as fixed factors, and with the inclusion of the interaction between pinnae and frequency. For both numerical and 3D models, we carried out post hoc tests between pinnae (y/n) at each frequency using estimated marginal means from the package *emmeans*.

**Supplementary Materials and Methods: Video recordings**

***V1: 3D print ear with microphone***

Video recording of probe microphone placement inside the 3D printed ear of *C. gorgonensis*. A digital micromanipulator with a holder restraining the 3D printed ear moved the ear along the probe tip. The microphone remained stationary. Scaled stimuli 6.67 kHz (60 kHz).

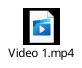

***V2: 3D print ear with microphone receiving broadband chirp***

3D printed ear of *C*. *gorgonensis* (1:11.512) receiving a scaled broadband chirp of 2.6 to 17 kHz (corresponding to 30 to 200 kHz) as the ear is moved into position with the probe microphone inside the cavity. Gain shown in magnitude (mPa).

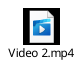

***V3: 3D print ear refractometry***

Quantitative imaging of acoustic waves using refracto-vibrometry in the field around the 3D printed ear of *C. gorgonensis* (Malkin et al., 2014). Screen recording software of scaled stimuli 9.63 kHz (110 kHz). Note the wave passing over the ear and the piston motion of the air inside showing the effect of the Helmholtz resonator.

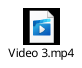
